# Functionalized nanoparticle transforms “cold” to “hot” adenoid cystic carcinoma of salivary gland tumour microenvironment in vitro

**DOI:** 10.64898/2026.04.18.719423

**Authors:** Rajdeep Chakraborty, Ria Shah, Arthur Chien, Masuma Akter, Ardeshir Amirkhani, Thiri Winn, Chao Shen, Mohammad-Ali Shahbazi, Anastasiia Tukova, Kerwin Shannon

## Abstract

Adenoid cystic carcinoma (ACC) of salivary gland is a “immune-cold” tumour. Annexin A3 (ANXA3) is an apoptotic protein found to be participating in immune cell infiltration in tumour microenvironment (TME) of various cancer cases. Significant low expressions of ANXA3 protein found in adenoid cystic carcinoma. We hypothesized overexpressing ANXA3 transforms ACC “cold” TME to “hot”. We cultured UM-HACC-2A and UFH2 spheroids on extracellular matrix and co cultured them with peripheral blood mononuclear cells. We functionalized FDA (The Food and Drug Administration) approved Poly(lactic-co-glycolic acid) PLGA nanoparticles with anti-cMyb antibody and ANXA3 recombinant protein using streptavidin-biotin conjugation. Upon overexpressing ANXA3 in ACC spheroids in immune coculture model using functionalized nanoparticles, significant increase of tumour infiltrating lymphocytes and decrease in the size of the ACC spheroids observed. Apoptotic profiler assay further confirmed significant upregulation of apoptotic proteins, some of them participate in immune infiltration. Overall, this project exhibits promising results showing potential approach to convert ACC into an immune “hot” tumour.

## INTRODUCTION

Treatment options for surgically inoperable or metastatic salivary gland cancer are restricted to chemotherapy. To date immunotherapy that includes immune check point inhibitors like Pembrolizumab and Nivolumab are not approved by FDA for the management of salivary gland cancer [1]. KEYNOTE-028 (NCT02054806) showed objective response rate of 12% with a 4-month median duration of response [1] and overall response rate of nivolumab was 4% [2]. Mucoepidermoid carcinoma (MEC) exhibit immune-hot tumour microenvironment (TME), whereas adenoid cystic carcinoma (ACC) shows contrasting features of immune-cold TME [3]. Objective response rate of immune check point inhibitors depends on the expression of PD-1 expression [3]. MEC express PD-1, showed an impressive objective response rate of approximately 33%, whereas ACC show negative PD-1 expression, presented a poor response rate of approximately 4-8% [3].

Immunological subtyping of salivary gland cancer identified variation of several pathways that includes apoptotic pathway in adenoid cystic carcinoma and mucoepidermoid carcinoma that might impact immunotherapy, due to diverse intrinsic cell compositions based on histologic origin [4]. So far, apoptosis has been found to tune tumour immunity based on the demonstration of apoptosis of immune cells (T effector cells), that undermines the anti-tumour reactivity of immune cells in the TME. This eventually facilitates the growth of T regulatory cells, M2 macrophage, and myeloid derived suppressor cells [5].

We previously found high expression of immunosuppressive proteins in salivary gland cancer cells that might be responsible for the lowering of immunotherapy efficacy [6]. Thus, based on the previous findings, we initially performed 3D-salivary gland – immune co-culture model construction and discovery phase mass spectrometry. We found significant low protein expression of Annexin A3 (ANXA3) in ACC compared to MEC cell lines. ANXA3 promotes immune infiltration and improves tumour prognosis [7], this might be one of the possible reasons for MEC exhibiting an immune-hot TME, whereas ACC shows features of immune-cold TME [3].

We, therefore, hypothesized that overexpression of Annexin A3 using functionalized nanoparticle will transform ACC into an immune hot tumour. To substantiate the hypothesis, we functionalized both gold nanoparticles (as a proof-of-concept) and PLGA nanoparticles (FDA approved) with anti-cMyb antibody. We did biophysical chemistry and in vitro characterization of the functionalized nanoparticles. After determining successful immobilization or conjugation of anti-cMyb antibody onto nanoparticles, we overexpressed Annexin A3 by conjugating recombinant ANXA3 onto anti-cMyb antibody functionalized PLGA nanoparticles. We determined the successful conjugation of recombinant ANXA3 protein using western blot and confocal microscope images. We overexpressed ANXA3 in ACC spheroids in PBMC coculture model and determined immune infiltration and size of the spheroids.

## RESULTS

### 3.1 Variations in protein expressions among ACC and other salivary gland pathoses

Upon SWATH analysis of MEC and ACC cell lines in immune coculture model, we found approx. 1000 proteins showing differential expressions. 2D cell culture and 3D cancer immune cell coculture models showed variation in the protein expressions (Figure 1a). Among them, the significantly upregulated and downregulated CD and apoptosis proteins are visualized using a heat map (Figure 1a). Annexin proteins and CD proteins showed significant variations among MEC and ACC cells in 3D coculture models (Figure 1a).

**Figure 1.**
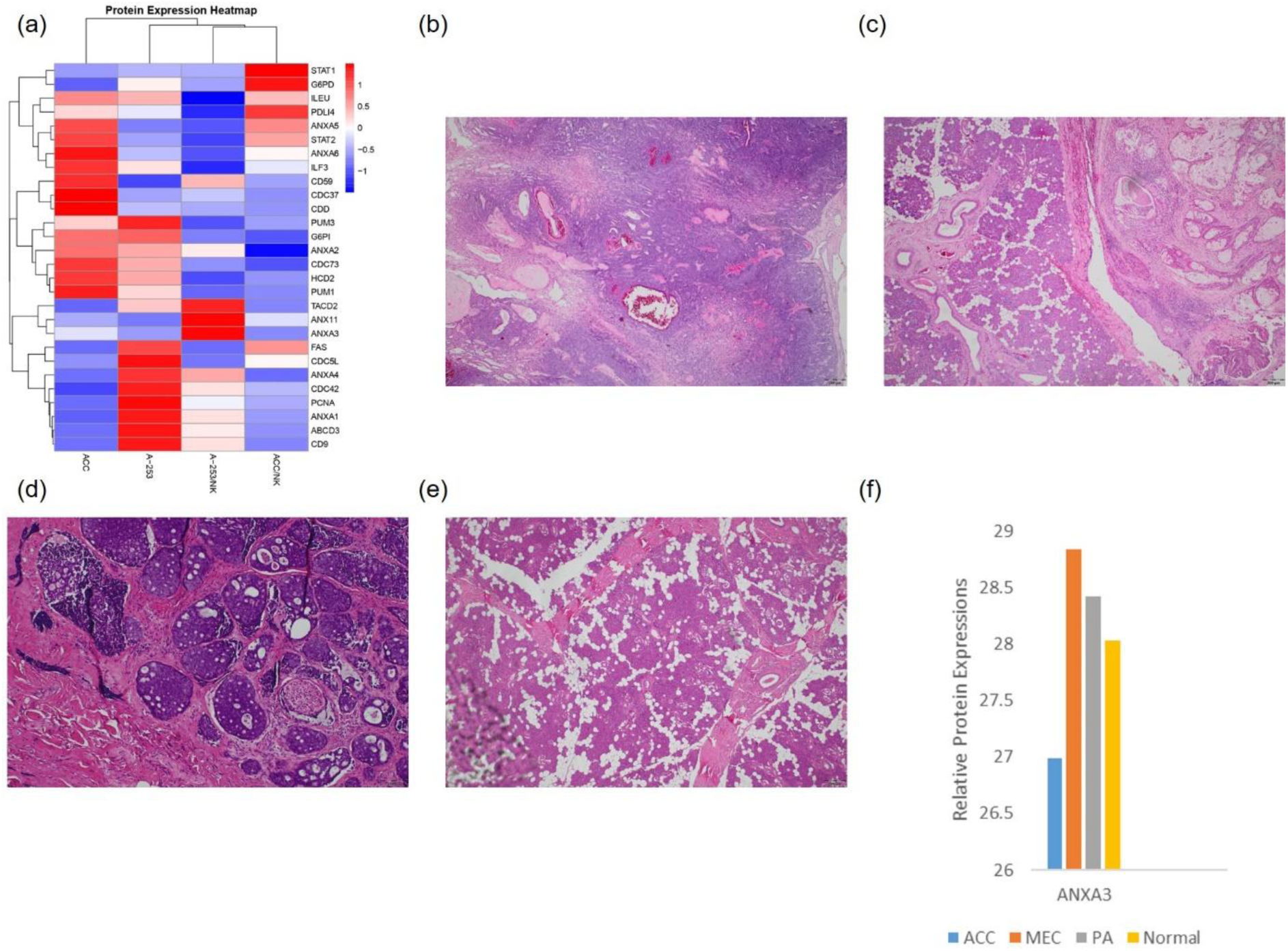
Mass Spectrometry Analysis of ACC cells and tissues. (a) SWATH data-based comparison of mechanism protein expressions among mucoepidermoid carcinoma and adenoid cystic carcinoma of salivary gland cancer cells in 2D and 3D cancer immune coculture model. Heat map to visualize hierarchical clustering of mechanism proteins in A-253 and ACC cells in 3D immune co-culture model made with heatmap (2) [gplots R package]. The protein intensities are log10 transformed and are displayed as colours ranging from blue to red as shown in the key. Both rows and columns are clustered using correlation distance and average linkage; Hematoxylin-eosin stained FFPE sample of (b) pleomorphic adenoma of salivary gland; Scale 200 µm; (c) mucoepidermoid carcinoma of salivary gland; Scale 200 µm; (d) adenoid cystic carcinoma of salivary gland; Scale 100 µm; (e) normal parotid salivary gland; Scale 200 µm; (f) bar chart showing ANXA3 relative protein expressions in FFPE samples.

**Figure 2.**
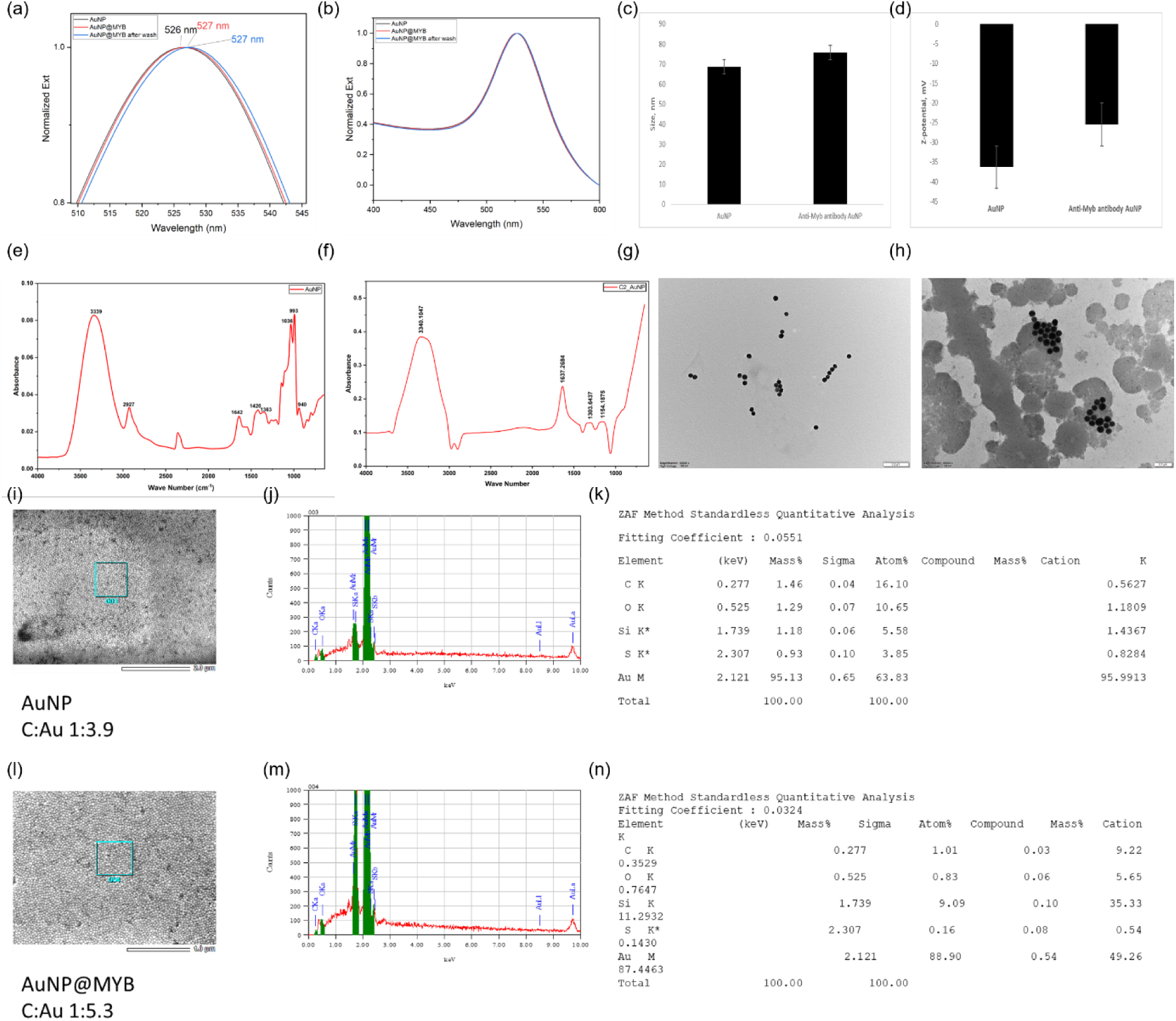
Biophysical chemistry characterization of anti-cMyb antibody-Au nanoparticles. (a) and (b) UV-Vis spectrum of the gold nanoparticles; (c) DLS; (d) Z-potential; (e) FT-IR of bare gold nanoparticles; (f) FT-IR of anti-cMyb antibody-Au nanoparticles; (g) TEM of bare gold nanoparticles; (h) TEM of anti-cMyb antibody-Au nanoparticles; (i). (j), (k) EDX analysis of bare gold nanoparticles; (l), (m), (n) EDX analysis of anti-cMyb antibody-Au nanoparticles. The error bar represents standard error of the mean. Red dots indicate regions of high Raman intensity at ∼1650 cm⁻¹, assigned to the amide I region (Figure 3a, b). This highlighted areas where the tissue interacts with the plasmonic surface of the AuNPs. In contrast, tissue regions not in contact with AuNPs do not exhibit measurable Raman signal intensity.

**Figure 3.**
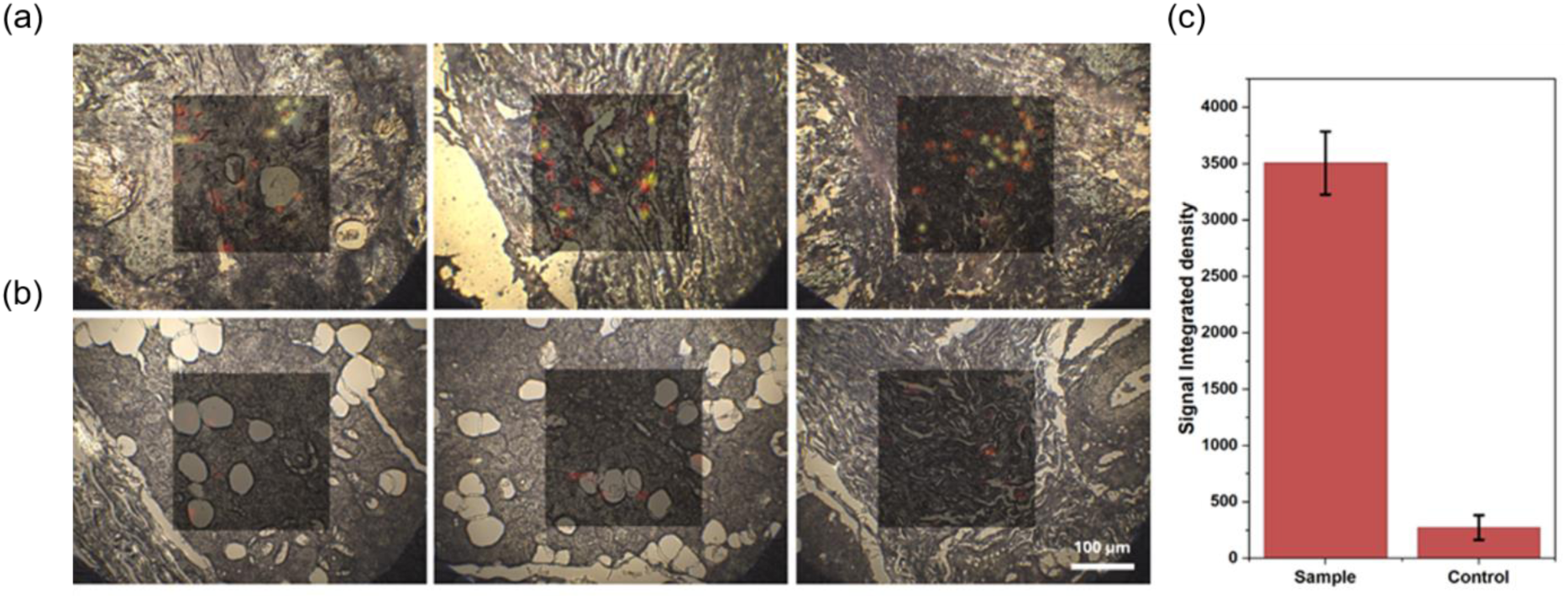
Raman 2D mapping of functionalized anti-cMyb antibody gold nanoparticles. (a) ACC; (b) normal parotid gland tissue samples; (c) bar chart showing comparison of signal integrated density of sample (ACC) and control (normal parotid gland). Scale 100 µm. The error bar represents standard error of the mean.

ANXA5 is significantly upregulated in ACC, whereas ANXA3 is significantly downregulated in ACC, which is opposite in MEC (Figure 1a). CD9, ANXA4, ANXA1, and ATP-binding cassette sub-family D member 3 are significantly high in MEC cells (Figure 1a). Additionally, G6PD, G6PI, FAS, ANXA6, CDC5L, CDC73, STAT1, and ILF3 expressions are high in both MEC and ACC, without showing any significant variations among the two salivary gland cancer cells (Figure 1a). Salivary gland cancer cells also showed presence of PCNA, CDC37, CDC42, STAT2, ANXA2, PUM3, PDL14, CDD, ILEU, and HCD2, but without significant variation among MEC and ACC (Figure 1a).

Upon hematoxylin-eosin stained histopathological section analysis, sample showed clear encapsulated/well cicumscribed tissue with finger like protrusion from the main mass. Tissue showed ductal, myoepithelial, and stromal components. Myoepithelial cells showed clear cytoplasm and many of the myoepithelial cells showed eccentric nuclei and eosinophilic cytoplasm resembling plasma cells. Multiple clear tyrosine crystals (amorphous eosinophilic floret like crystals) and epithelial sheets and spindle-shaped mesenchymal cells presence confirmed the sample to be pleomorphic adenoma of salivary gland (Figure 1b).

Other sample showed tissue composed of solid and cystic areas. Epidermoid, mucous, and intermediate cells observed, where the squamous component show distinct cellular boundaries. Focal areas of clear cell changes observed. Sheets of neoplastic squamoid cells with crowded and overlapping nuclei, and mucous cells seen in clusters. Multiple groups of keratinizing squamous neoplastic cells observed. Thus, the sample confirmed to be mucoepidermoid carcinoma of salivary gland (Figure 1c).

Another sample showed histopathological features of tubular pattern composed of inner ductal and outer myoepithelial cells, where the ductal cells are cuboidal with eosinophilic cytoplasm. Myoepithelial cells form cribriform pattern, additionally sample showed comedo type tumour necrosis, frequesnt mitosis, and marked nuclear atypia with prominent central nucleoli. Frequest perineural and intraneural invasions observed. Taken together, the sample is confirmed to be adenoid cystic carcinoma of salivary gland (Figure 1d).

The FFPE slide showed histologic characters like serous acini, intercalated ducts, and intervening adipose tissue. The serous acini cells are polygonal shaped with dense, intracytoplasmic, basphilic zymogen granules. The intercalated ducts are lined by single layer of cuboidal epithelium. Striated ducts are lined by simple columnar epithelium consisting densely packed mitochondria that form cytoplasmic striations. The interlobular ducts are observed to be located within interlobular septa and lined by pseudostratified columnar epthelium. Possible sebacous cells found within the wall of striated duct and the sample additionaly showed the presence of intraparenchymal lymph nodes. The sample confirmed to be normal parotid gland (Figure 1e).

Upon SWATH analysis of FFPE samples of normal parotid salivary gland, pleomorphic adenoma of salivary gland, mucoepidermoid carcinoma of salivary gland, and adenoid cystic carcinoma, ANXA3 expression found lowest in ACC and highest in MEC samples (Figure 1f).

### 3.2 Functionalised anti-cMyb antibody functionalised nanoparticles

Since cMyb is overexpressed in ACC [8], to target ACC, first anti-cMyb antibody was conjugated to nanoparticles. To acquire a proof-of-concept data showing successful immobilization of anti-cMyb antibody on nanoparticles, we initially used gold nanoparticles. The UV-Vis spectrum of the gold nanoparticles showed a surface resonance peak at 526 nm, indicating a size range of approximately 30-50 nm. Which is in accordance to the supplier product description of gold nanoparticles size 40 nm. Upon addition of anti-cMyb antibody, a reproducible, albeit small, 1 nm red shift 527 nm observed. The Z-average hydrodynamic diameter of the nanoparticles was found to be approximately 73.6 nm and z-potential approximately -36.7 mV. Upon addition of anti-cMyb antibody, the Z-average diameter increased by approximately 1 nm and z-potential increased to approximately -20.5 mV. It indicates that the anti-cMyb antibody were successfully conjugated without causing significant aggregation, forming a relatively dense, mono-layered, or flat-oriented corona rather than a thick, multi-layered or aggregated structure.

Upon FTIR spectra analysis, peptide linkage identified – Amide I 1637 cm^-1^ sharp peak confirms conjugation. The sharp peak is primarily due to C=O with minor C-N stretching. Broad peak at 3340 cm^-1^ denotes O-H stretching and N-H stretching characteristic of amide bonds (-NH), hydroxyl groups (-OH), amino groups (-NH_2_) present in the amino acid backbone and side chains of the antibody. This broad peak after gold-antibody conjugation, is primarily attributed to the presence of protein layers on the surface of gold nanoparticles. FTIR showing sharp peak at 2750-3000 cm^-1^ is characteristic of C-H stretching vibrations that confirms antibody attached to surface of gold nanoparticles. Sharp peak at 2250 cm^-1^ denotes successful covalent bonding reaction (antibody linked to a spacer on the nanoparticle). Peak at 3750-4000 cm^-1^ denotes presence of structural elements with N-H and O-H vibrations that are not involved in strong hydrogen bonding, that is, presence of “free” or “weak” bonded groups in vibrational spectroscopy. Finally, peak at 1154 cm^-1^ denotes presence of capping agents; organic capping shell prevents gold nanoparticles from aggregation. The thiol group (-SH) in the molecule forms a strong covalent bond with the gold surface (Au-S), while the hydrophilic polyethylene glycol (PEG) chain provides steric repulsion, preventing the particles from sticking together.

High-resolution TEM confirms spherical structure of gold nanoparticles without aggregation, with an average size of 22.537 nm. The size of the gold nanoparticles increased significantly upon conjugation with anti-cMyb antibody, with an average size of 32.48 nm, that suggests formation of an anti-cMyb antibody corona layer.

Energy Dispersive X-ray Spectroscopy (EDX) showed an increased carbon-to-gold (C/Au) ratio upon antibody conjugation to gold nanoparticles (1:5.3 vs 1:3.9), and an increase of approximately 55% of atomic percentage of oxygen compared to non-conjugated gold nanoparticle, further confirms successful bioconjugation of anti-cMyb antibody to gold nanoparticles.

Upon acquiring proof-of-concept preliminary data showing successful conjugation of anti-cMyb antibody onto gold nanoparticle, we further proceed to using FDA (The Food and Drug Administration) approved Poly(lactic-co-glycolic acid) PLGA nanoparticles for conjugating anti-cMyb antibody, overexpressing ANXA3 protein and adding the functionalized nanoparticles to ACC spheroid immune coculture model to determine whether ANXA3 overexpression leads to increase in infiltration of tumour infiltrating lymphocytes.

Streptavidin coated PLGA nanoparticles were added to biotinylated anti-cMyb antibody and biophysical chemistry assays were performed. The UV-Vis spectrum of the PLGA nanoparticles showed an absorption band at 272 nm, indicating a size range of approximately 120-300 nm. Which is in accordance to the supplied product description of PLGA nanoparticles size 120 nm. Upon addition of anti-cMyb antibody, 2 nm red shift observed. The Z-average hydrodynamic diameter of the nanoparticles was found to be approximately 408 nm and z-potential approximately -12 mV. Upon addition of anti-cMyb antibody, the Z-average diameter increased by approximately 10 nm and z-potential increased to approximately -8 mV. It indicates that the anti-cMyb antibody were successfully conjugated onto PLGA nanoparticles.

A sharp drop or disappearance of peaks in the 1000 cm^-1^ to 1100 cm^-1^ range in FTIR spectra after adding biotinylated antibody to PLGA-streptavidin coated nanoparticles is a strong indication of successful surface conjugation. This region (represent C–O stretching) typically corresponds to vibrational modes that become hidden or “screened” when the streptavidin-coated particle surface binds to the biotinylated antibody, effectively coating the previously exposed surface and changing its absorption profile. Peptide linkage identified – Amide I 1624 cm^-1^ sharp peak confirms conjugation. The sharp peak is primarily due to C=O stretching. N-H bending and C-N stretching of the amide bond (amide II) observed at 1575 cm^-1^ of the spectra. Broad band from 3383 cm^-1^ indicating N–H stretching (amide A) and OH stretching from the antibody.

High-resolution TEM confirms uniform, spherical structure with distinct, well-defined boundaries of PLGA nanoparticles, with an average size of 127.165 nm. The size of the PLGA nanoparticles exhibits a larger diameter or a “shell” upon conjugation with anti-cMyb antibody, with an average size of 129.874 nm, that suggests formation of an anti-cMyb antibody corona layer.

EDX spectroscopy results showed an increased atomic concentration of carbon, and a three-fold increase in oxygen atomic concentration upon anti-cMyb antibody conjugation to PLGA nanoparticles. EDX elemental mapping detects the emergence of nitrogen (atomic concentration of 15.67) and sulfur (atomic concentration of 3.15) peaks on the nanoparticle surface, which are absent in pure PLGA (Figure 4). The appearance of a distinct nitrogen peak is a direct indicator of antibody conjugation. Nitrogen originates from the amide bonds (peptide backbone) of the antibody (e.g., IgG, Fab fragments). The sulfur peak is attributed to sulfur-containing amino acids (cysteine and methionine) within the antibody’s structure. Thus, EDX spectroscopy further confirms successful bioconjugation of anti-cMyb antibody to PLGA nanoparticles.

**Figure 4.**
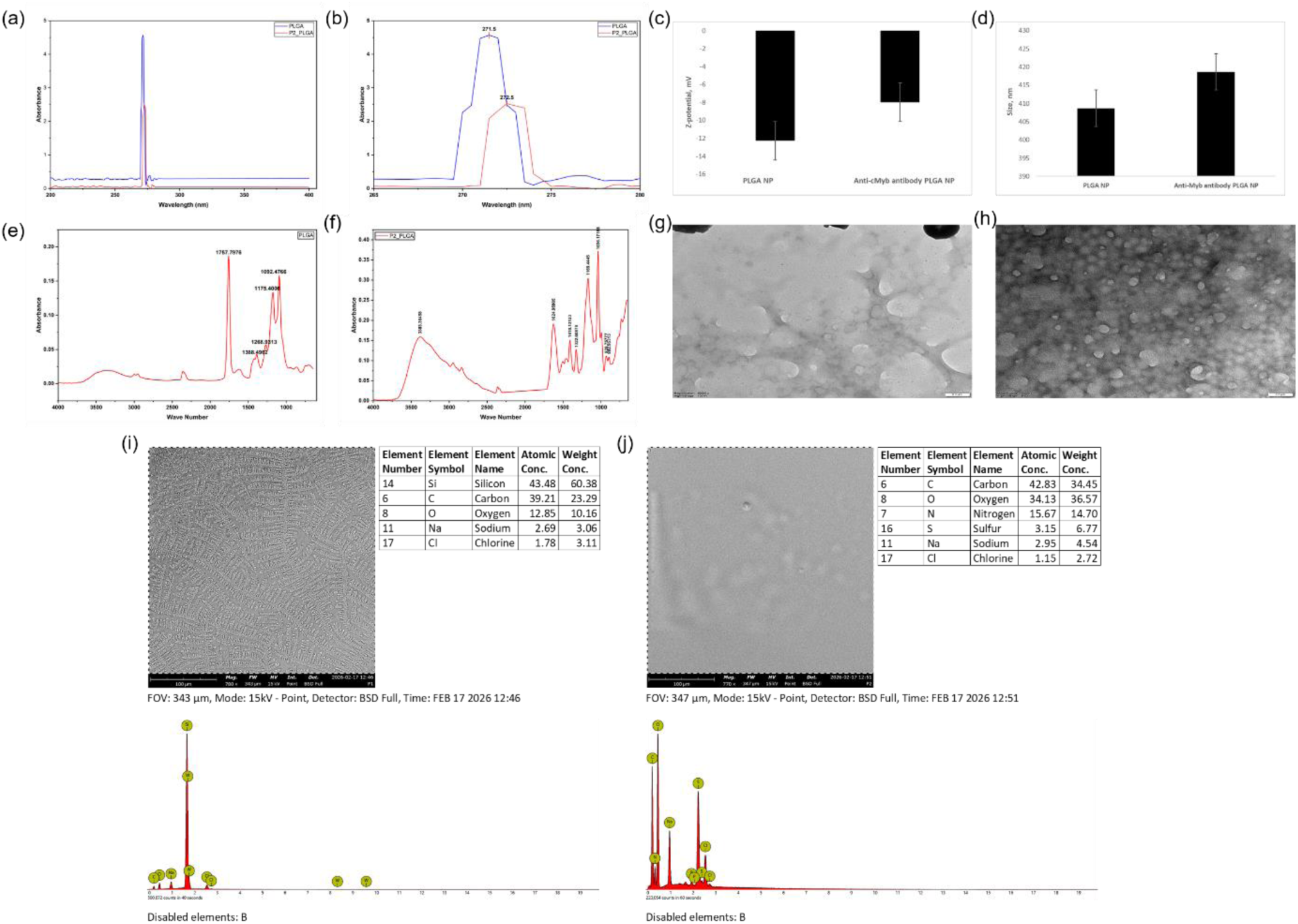
Biophysical chemistry characterization of anti-cMyb antibody-PLGA nanoparticles. (a) and (b) UV-Vis spectrum of the PLGA nanoparticles; (c) DLS; (d) Z-potential; (e) FT-IR of bare PLGA nanoparticles; (f) FT-IR of anti-cMyb antibody-PLGA nanoparticles; (g) TEM of bare PLGA nanoparticles; (h) TEM of anti-cMyb antibody-PLGA nanoparticles; (i). (j), (k) EDX analysis of bare PLGA nanoparticles; (l), (m), (n) EDX analysis of anti-cMyb antibody-PLGA nanoparticles. The error bar represents standard error of the mean.

Further, we completed *in vitro* characterization of the anti-cMyb antibody conjugated PLGA nanoparticles using salivary gland adenoid cystic carcinoma FFPE samples, normal salivary gland FFPE samples, UM-HACC-2A cells, UFH2 cells, and SCC9 cells (negative control) (Figure 5). Upon addition of biotinylated anti-Myb antibody conjugated streptavidin coated PLGA nanoparticles or otherwise anti-cMyb antibody functionalized PLGA nanoparticles showed highly specific opaque green fluorescent specs on the cribriform histologic patters of ACC tissue. Upon immunohistochemical staining with anti-cMyb antibody, abundant non-specific staining artefacts observed. Additionally, upon adding anti-cMyb antibody functionalized PLGA nanoparticles and anti-cMyb antibody, both showed no positive fluorescence specs in normal parotid gland FFPE sample. Further, Alexa Fluor 647 labelled anti-cMyb antibody functionalized PLGA nanoparticles were added to UM-HACC-2A cells, UFH2 cells, and SCC9 cells. Both UM-HACC-2A, UFH2 nucleus showed positive far-red Alexa Fluor 647 bright opaque fluorescent specs that denotes successful cMyb antigen-antibody reaction, whereas SCC9, which is a cMyb negative cell line, did not show any positive far-red Alexa Fluor 647 bright opaque fluorescent specs (Figure 5). Taking together the biophysical chemistry and *in vitro* characterization results, it was confirmed that anti-cMyb antibody functionalized PLGA nanoparticles are functional and can be used to target cMyb positive tissues and cells in further experiments.

**Figure 5.**
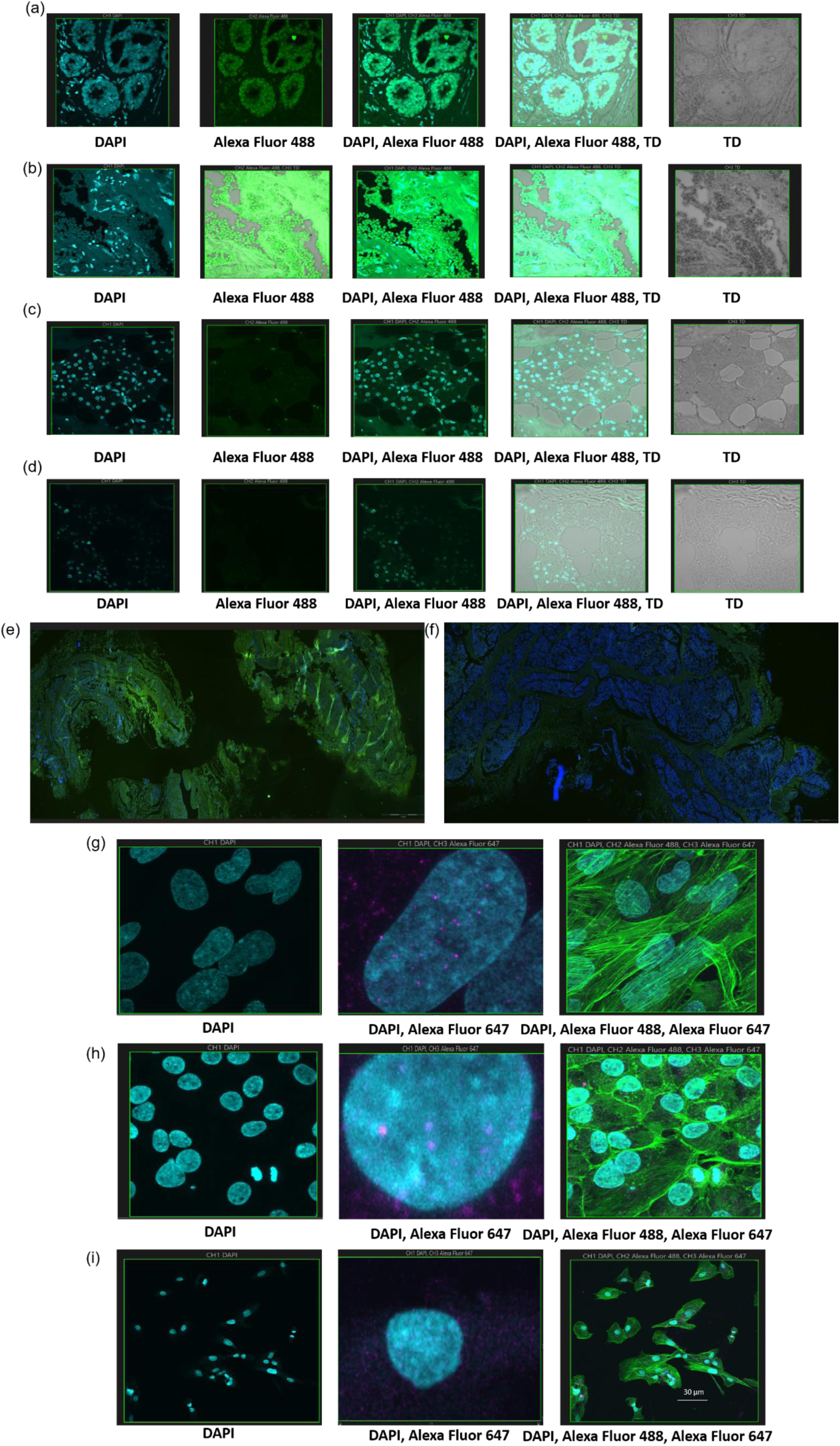
In Vitro FFPE and cell line characterization of anti-cMyb antibody-PLGA nanoparticles. (a) anti-cMyb antibody-PLGA nanoparticles on ACC; (b) anti-cMyb antibody on ACC; (c) anti-cMyb antibody-PLGA nanoparticles on normal parotid salivary gland; (d) anti-cMyb antibody on normal parotid salivary gland; widefield microscope images with 4X objective of (e) anti-cMyb antibody-PLGA nanoparticles on ACC; (f) anti-cMyb antibody-PLGA nanoparticles on normal parotid salivary gland; scale 200 µm; 3D z-stack confocal images with 60X objective of (g) anti-cMyb antibody-PLGA nanoparticles on UM-HACC-2A cells; (h) anti-cMyb antibody-PLGA nanoparticles on UFH2 cells; (i) anti-cMyb antibody-PLGA nanoparticles on SCC9 cells. Scale 30 µm.

### 3.3 Overexpression of ANXA3 protein

To substantiate our hypothesis, next, streptavidin coated PLGA nanoparticles were conjugated with biotinylated [Alexa Fluor 647 labelled anti-MYB antibody and recombinant ANXA3 protein (1:1)], and added to ACC cells to overexpress ANXA3 and assess immune infiltration in the 3D cell culture matrix. 3D z stack imaging confirmed significant amount of positive far-red Alexa Fluor 647 bright opaque fluorescent specs signifying successful uptake of ANXA3 conjugated anti-cMyb antibody functionalized PLGA nanoparticles in UM-HACC-2A and UFH2 cells. Whereas, upon addition of Alexa Fluor 647 labelled streptavidin PLGA nanoparticles did not show significantly observable fluorescence specs. Here, Alexa Fluor 647 labelled streptavidin PLGA nanoparticles acted as a negative control. Western blot analysis done to further confirm successful uptake of ANXA3 conjugated anti-cMyb antibody functionalized PLGA nanoparticles. ANXA3 Recombinant Protein acted as positive control, PLGA acted as negative control, UM-HACC-2A, UFH2 acted as experimental control, and GAPDH as internal control of the experiment. Western blot further confirmed successful uptake of ANXA3 conjugated anti-cMyb antibody functionalized PLGA nanoparticles in ACC cells, showing significant increase of ANXA3 protein (35 kDa) after using functionalized nanoparticles (Figure 6).

**Figure 6.**
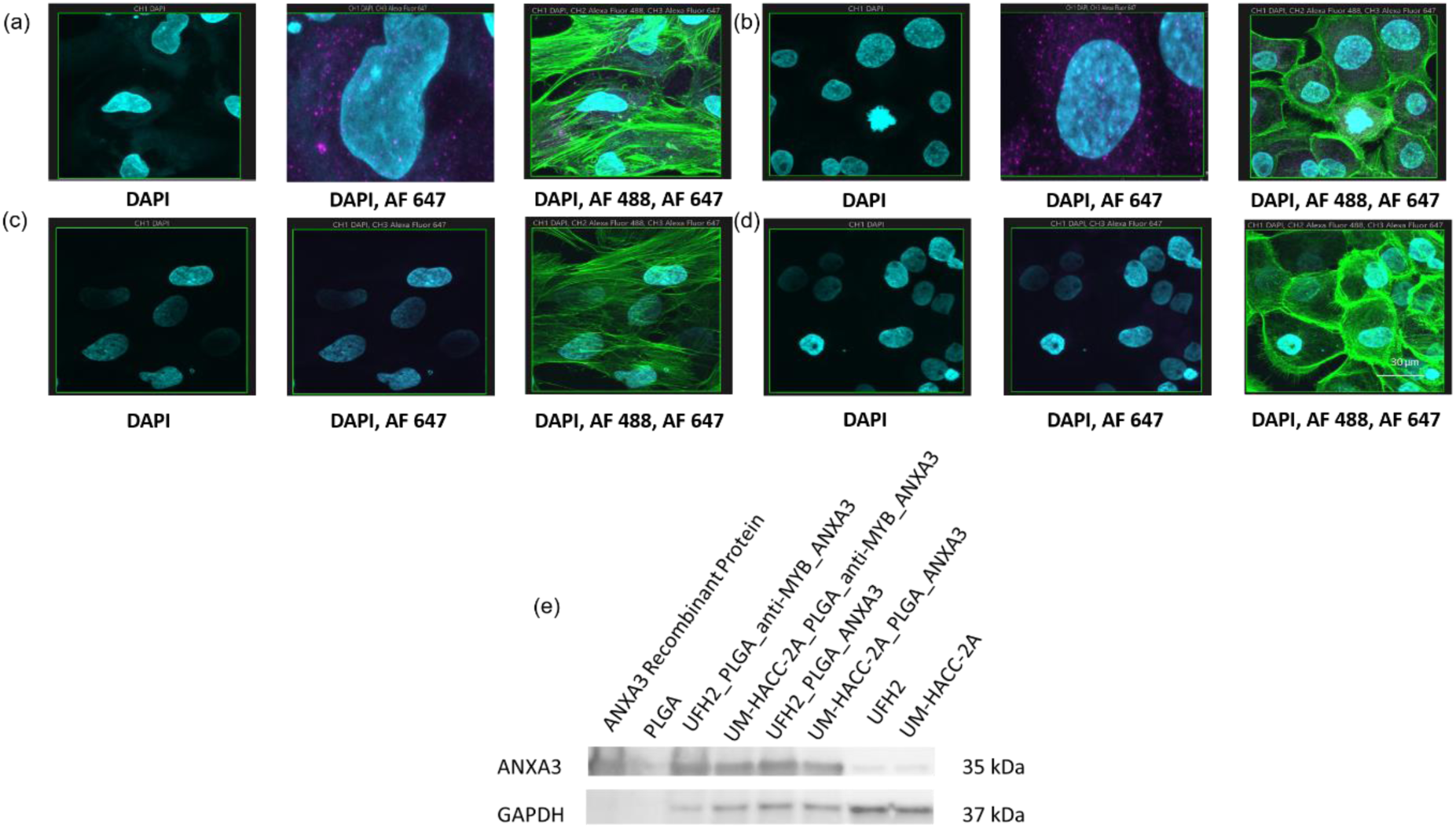
Functionalization of anti-cMyb antibody-PLGA nanoparticles with recombinant ANXA3 protein. Alexa Fluor 647 was conjugated to the biotinylated mixture using Alexa Fluor 647 conjugation kit (Fast) – Lightening Link (AB269823-1004, abcam). 3D z-stack confocal images with 60X objective of (a) anti-cMyb antibody-ANXA3-PLGA nanoparticles on UM-HACC-2A cells; (b) anti-cMyb antibody-ANXA3-PLGA nanoparticles on UFH2 cells; (c) Alexa Fluor 647-PLGA nanoparticles on UM-HACC-2A cells; (d) Alexa Fluor 647-PLGA nanoparticles on UFH2 cells; Scale 30 µm; (e) western blot showing overexpression of ANXA3 using anti-cMyb antibody-ANXA3-PLGA nanoparticles. UM-HACC 2A and UFH2 acted as a control, ANXA3 recombinant protein acted as a positive control and PLGA acted as a negative control. Positive control 4 µg, control 10 µg, and the rest of the samples 8 µg was loaded. Pre-stained protein ladder (ab116029, abcam), Rabbit annexin A3 polyclonal antibody (PA5-82483, Thermo Fisher Scientific) at 0.4 µg/mL, Goat anti-Rabbit IgG (H+L) Secondary Antibody, HRP (31460, Thermo Fisher Scientific) at 1:10,000, mouse GAPDH monoclonal antibody ZG003 (398600, Thermo Fisher Scientific) at 2 µg/mL, and Goat anti mouse IgG HRP (31430, Thermo Fisher Scientific) at 1:5000.

### 3.4 Overexpression of ANXA3 protein resulted in increased immune infiltration and reduction of the shape of the ACC spheroids

First, UM-HACC-2A and UFH2 spheroids stability was assessed on 3D ECM in a PBMC coculture model. Viability of UM-HACC-2A and UFH2 cells remained constant on ECM. UM-HACC-2A and UFH2 spheroids showed stable phenotype on 3D ECM and on 3D ECM in a PBMC coculture.

Upon addition of ANXA3 conjugated anti-cMyb antibody functionalized PLGA nanoparticles to UM-HACC-2A and UFH2 spheroids on 3D ECM in a PBMC coculture model resulted in significant increase of CD4+ and CD8+ positive lymphocytes in the ECM incrementally over time, that is, at 72 hours showed maximum infiltration of tumour infiltrating lymphocytes (Figure 7). The control groups UM-HACC-2A and UFH2 spheroids on 3D ECM in a PBMC coculture did not show much variation in CD4+ and CD8+ positive lymphocytes over time (Figure 8). Recombinant GAPDH protein (non-specific control) overexpression following similar protocol was done. The non-specific control did not result in significant change in CD4+ and CD8+ positive lymphocytes in ECM over time (Figure 8).

**Figure 7.**
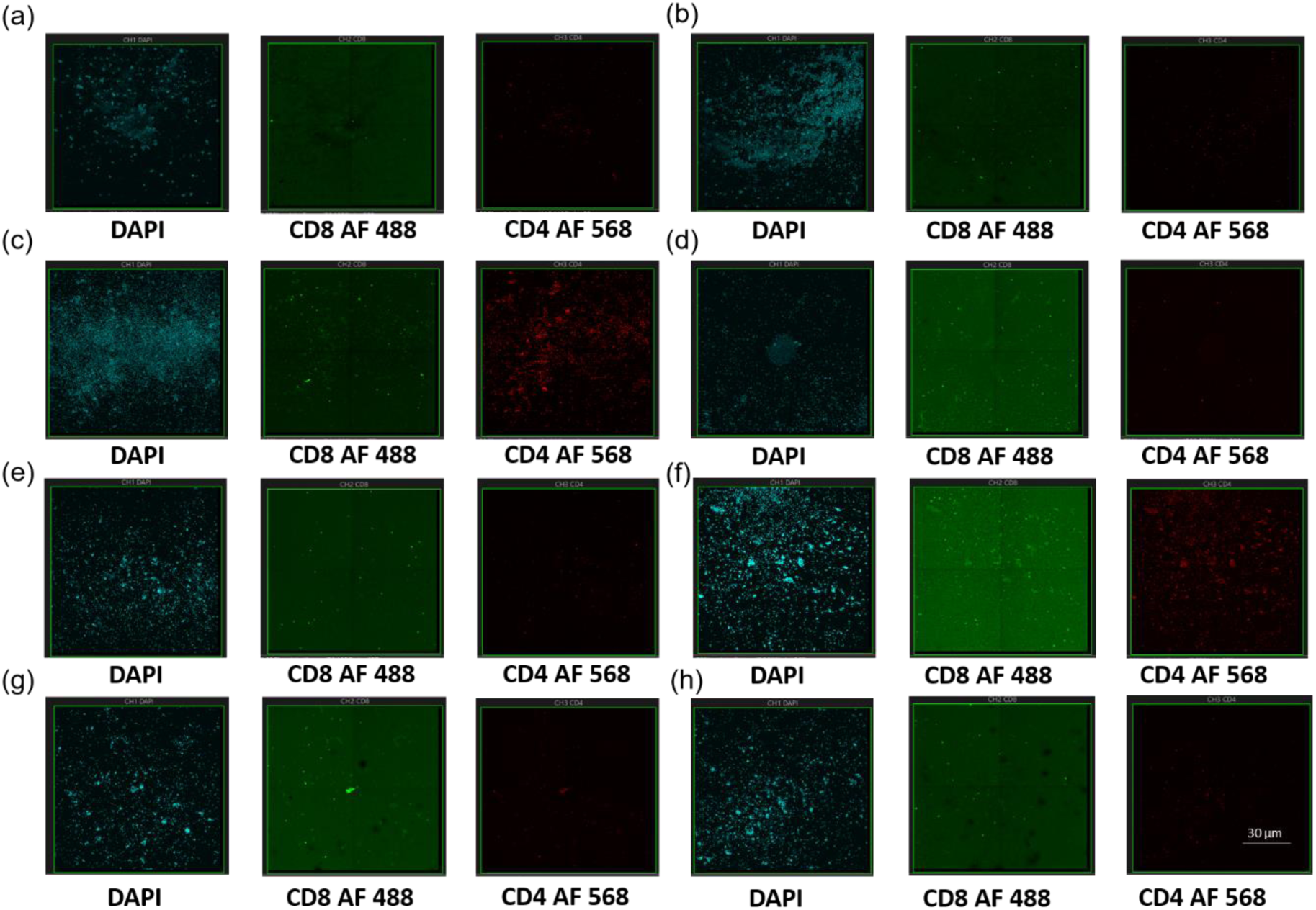
Infiltration of immune cells in ACC TME due to ANXA3. 3D z-stack confocal images of CD4+ and CD8+ with 60X objective of (a) anti-cMyb antibody-ANXA3-PLGA nanoparticles on UM-HACC-2A spheroids in PBMC coculture in 24 hours; (b) anti-cMyb antibody-ANXA3-PLGA nanoparticles on UM-HACC-2A spheroids in PBMC coculture in 48 hours; (c) anti-cMyb antibody-ANXA3-PLGA nanoparticles on UM-HACC-2A spheroids in PBMC coculture in 72 hours; (d) anti-cMyb antibody-ANXA3-PLGA nanoparticles on UFH2 spheroids in PBMC coculture in 24 hours; (e) anti-cMyb antibody-ANXA3-PLGA nanoparticles on UFH2 spheroids in PBMC coculture in 48 hours; (f) anti-cMyb antibody-ANXA3-PLGA nanoparticles on UFH2 spheroids in PBMC coculture in 72 hours; (g) anti-cMyb antibody-GAPDH-PLGA nanoparticles on UM-HACC-2A spheroids in PBMC coculture in 72 hours; (h) anti-cMyb antibody-GAPDH-PLGA nanoparticles on UFH2 spheroids in PBMC coculture in 72 hours. Scale 30 µm.

**Figure 8.**
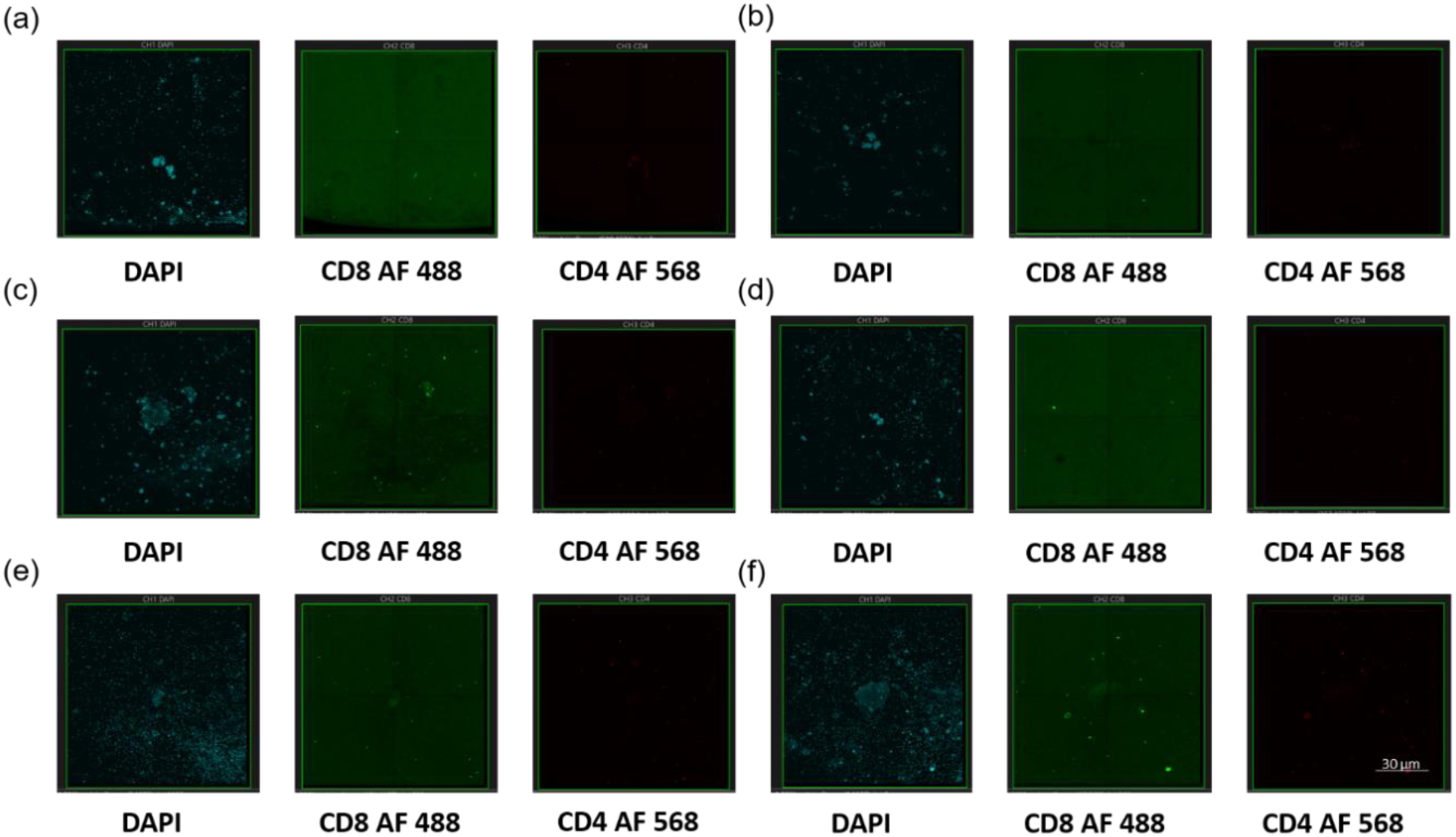
Infiltration of immune cells in ACC TME. 3D z-stack confocal images of CD4+ and CD8+ with 60X objective of (a) UM-HACC-2A spheroids in PBMC coculture in 24 hours; (b) UM-HACC-2A spheroids in PBMC coculture in 48 hours; (c) UM-HACC-2A spheroids in PBMC coculture in 72 hours; (d) UFH2 spheroids in PBMC coculture in 24 hours; (e) UFH2 spheroids in PBMC coculture in 48 hours; (f) UFH2 spheroids in PBMC coculture in 72 hours. Scale 30 µm.

Ostensibly, the UM-HACC-2A and UFH2 spheroids showed significant reduction in size upon addition of ANXA3 conjugated anti-cMyb antibody functionalized PLGA nanoparticles in the PBMC coculture model compared to the control groups (Figure 9).

**Figure 9.**
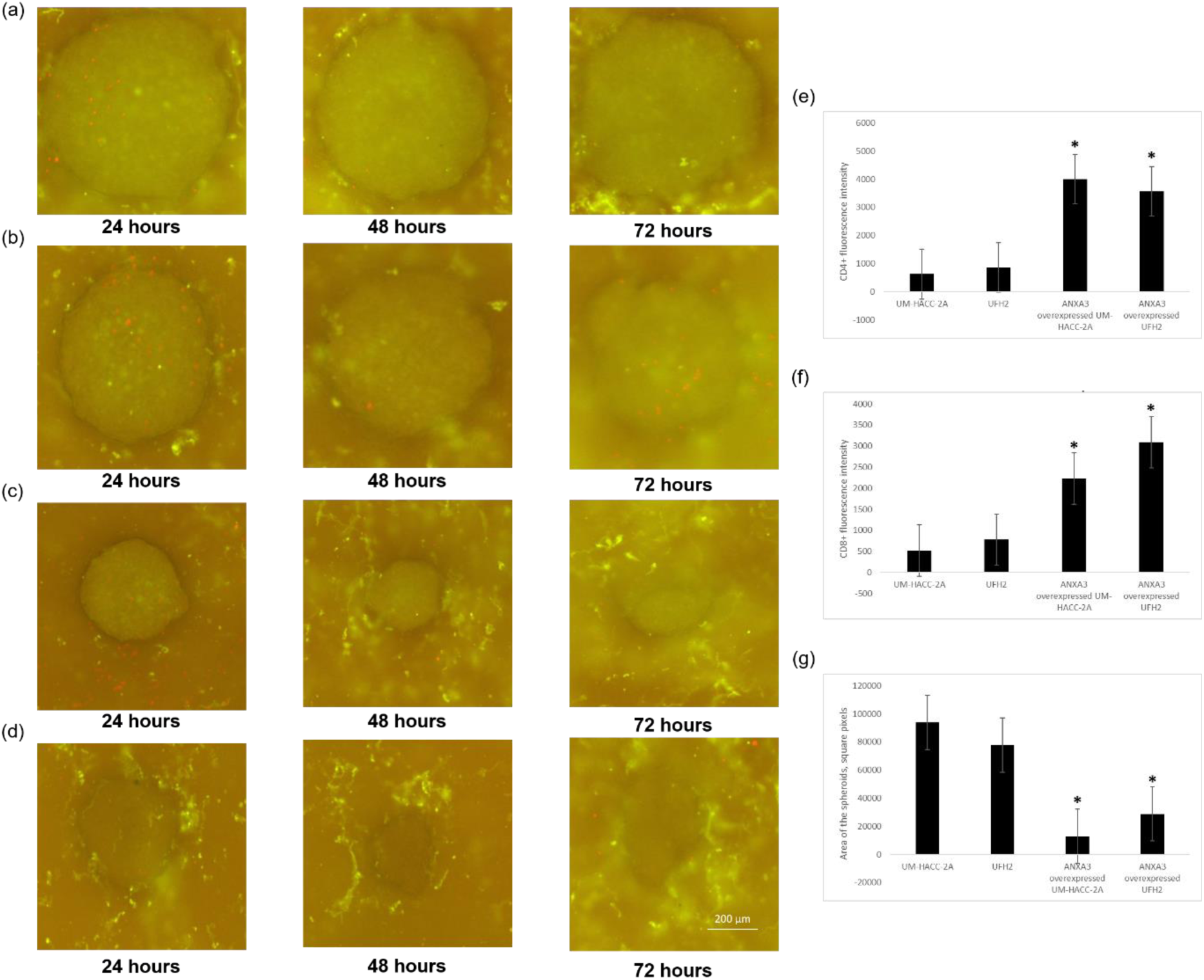
Effect of immune cells infiltration on ACC spheroids due to ANXA3. Widefield microscope images with 4X objective of (a) UM-HACC-2A spheroids in PBMC coculture; (b) UFH2 spheroids in PBMC coculture; (c) anti-cMyb antibody-ANXA3-PLGA nanoparticles on UM-HACC-2A spheroids in PBMC coculture; (d) anti-cMyb antibody-ANXA3-PLGA nanoparticles on UFH2 spheroids in PBMC coculture. Scale 200 µm. (e) CD4+ fluorescence intensity of UM-HACC-2A spheroids in PBMC coculture, UFH2 spheroids in PBMC coculture, anti-cMyb antibody-ANXA3-PLGA nanoparticles on UM-HACC-2A spheroids in PBMC coculture, and anti-cMyb antibody-ANXA3-PLGA nanoparticles on UFH2 spheroids in PBMC coculture (n=6); \**P*≤0.05. The error bar represents standard error of the mean.

### 3.5 Overexpression of ANXA3 protein resulted in variation in apoprotic proteins

Upon apoptotic profiler result analysis, Pro caspase 3 and cleaved caspase 3 expressions were found significantly higher in adenoid cystic carcinoma spheroids treated with ANXA3 conjugated anti-cMyb antibody functionalized PLGA nanoparticles (Figure 10).

**Figure 10.**
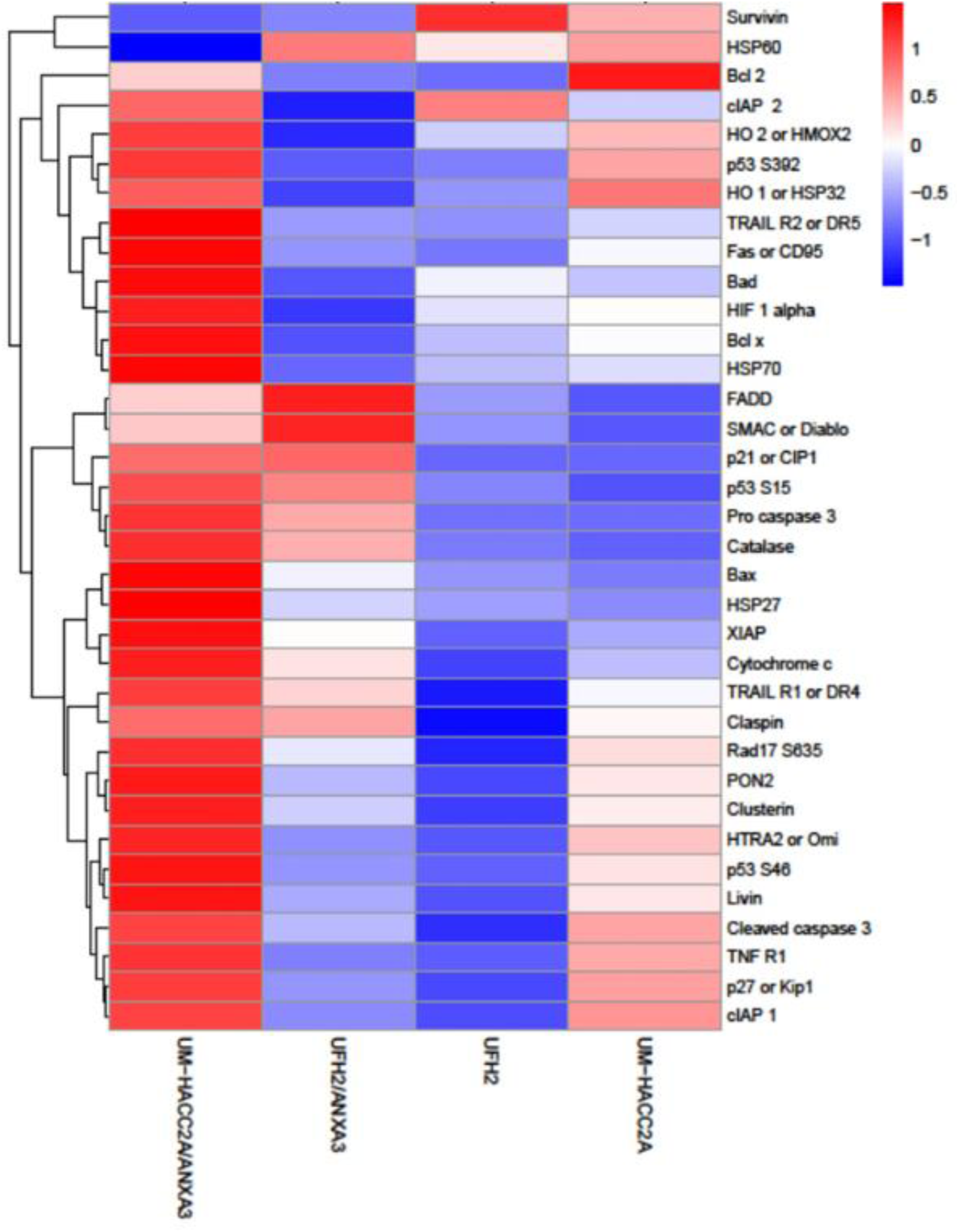
Differential apoptotic proteins due to the effect of immune infiltration induced by ANXA3 in ACC spheroids. Comparison of apoptotic protein expressions in UFH2 in PBMC coculture, UM-HACC-2A in PBMC coculture, anti-cMyb antibody-ANXA3-PLGA nanoparticles on UM-HACC-2A spheroids in PBMC coculture, and anti-cMyb antibody-ANXA3-PLGA nanoparticles on UFH2 spheroids in PBMC coculture. Heat map to visualize hierarchical clustering of apoptotic proteins in A-253 and ACC cells in 3D immune co-culture model made with heatmap.2() [gplots R package]. The protein intensities are log10 transformed and are displayed as colours ranging from blue to red as shown in the key. Both rows and columns are clustered using correlation distance and average linkage.

Additionally, HSP27, SMAC, and catalase are significantly high in functionalized nanoparticle treated ACC spheroids to untreated (Figure 10). Ostensibly, survivin are significantly lower in

ANXA3 overexpressed ACC spheroids. Fas-Associated Death Domain (FADD) is significantly lower in the same compared to the control (Figure 10). Interestingly, Bad, Bcl x, HIF 1 alpha, HSP60, and HSP70 showed variation in expression changes in UM-HACC-2A and UFH2 spheroids after ANXA3 overexpression (Figure 10).

p53 at position serine 392, 46, and 15 expressions are significantly high in ANXA3 overexpressed ACC spheroids (Figure 10). Similarly, PON2, TRAIL R1 or DR4, p21 or CIP1, TRAIL R2 or DR5, Bax, p27 or Kip1, Bcl 2, Fas or CD95, HO 1 or HSP32, Rad17 S635, HO 2 or HMOX2, cIAP 1, cIAP 2, TNF R1, Claspin, XIAP, HTRA2 or Omi, Clusterin, Livin, and Cytochrome c showed significant higher expressions in ANXA3 overexpressed ACC spheroids (Figure 10).

## Discussion

Cancer cells launch counter-assaults against anti-tumour reactivity of immune cells by its intrinsic apoptosis machinery that incorporate activation of both extrinsic and intrinsic apoptosis related proteins. ANXA3, which is an apoptotic protein found to be involved in immune infiltration [9]. High ANXA3 expressions correlates with a favourable prognosis of ovarian serous carcinoma [7]. ANXA3 increases T follicular helper cells and T lymphocytes infiltration in the tumour microenvironment of ovarian serous carcinoma [7]. Contrastingly, ANXA3 promotes tumorigenicity in hepatocellular carcinoma by modifying the immune microenvironment by increasing neutrophil-lymphocyte ratio and decreasing T-lymphocyte in the tumour microenvironment [9]. Similarly, in the laryngeal squamous cell carcinoma, ANXA3 promotes M2 (tumour associated macrophage), creating an immunosuppressive tumour microenvironment [10]. Apart from cancer cases, ANXA3 exhibited leukocyte infiltration and inflammatory responses in sepsis [11], acting as a potential biomarker for diagnosis. Similar to the previous observations, upon overexpressing ANXA3 in ACC spheroids, tumour infiltrating lymphocytes infiltration increased. This result is again in accordance with the previous findings that suggest high expression of ANXA3 promotes immune infiltration and improves tumour prognosis [7].

Increased infiltration of tumour infiltrating lymphocytes generally correlates with smaller tumour size, improved clinical outcomes, and better response to immunotherapy [12]. Triple-Negative Breast Cancer (TNBC) and HER2-positive breast cancer, high TILs are associated with smaller tumor diameter and lower histological grade [13]. Infiltrated CD8+ T cells release cytotoxic granules (perforin/granzyme) or use Fas/FasL system to induce apoptosis in cancer cells [14]. In our study, we found the same, ACC spheroid size significantly reduced upon ANXA3 overexpression, that lead to the determination of the variation of apoptotic proteins in ACC spheroids induced by increased immune cell infiltration.

The most significant and well-studied immune suppressive protein related to apoptosis is p53. Interestingly, p53 abrogation contribute to tumour immune evasion. The p53 mutation regulates major histocompatibility complex molecules that eventually diminishes cancer cell capability to provoke immune response [15]. Phosphorylated p53 expressions are observed in both ACC and MEC cells during this project. Significant variation of p53 found in ACC spheroids due to overexpression of ANXA3 protein.

Pro-caspase 3 expressions are significantly high in ANXA3 overexpressed ACC spheroids. Previously, targeted therapies, such as BH3-mimetics and molecular inhibitors, have been shown to induce apoptosis in ACC cells by triggering caspase activity [16].

Similarly, Survivin which is significantly lower in ANXA3 overexpressed ACC, is an inhibitor of apoptosis and also related to poor prognosis. Survivin is known to trigger immune effector response. Although, relative contribution of various splice variants of Survivin in immune evasion is still elusive [17]. Significant lower expression of FADD observed in ANXA3 overexpressed ACC compared to control. Higher expression of FADD is frequently linked to “immunologically cold” tumour characterized by high infiltration of suppressive immune cells (T regulator and M2 cells). Low expression of FADD often correlates with higher infiltration of CD8+ T cells, plasma cells, and memory B cells [18].

HMOX1, Bad, Bax, Bcl-2, and CD95 expressions are found significant higher expressions in ANXA3 overexpressed ACC. Paradoxically, CD95 participate in immune suppressive mechanisms [19]. Our data suggest that both HSP27 and catalase found significantly high in ANXA3 overexpressed ACC spheroids. HSP27 is a pro-apoptotic protein [20], whereas catalase has been implicated as anti-apoptotic [21]. Additionally, Catalase has been found to be playing critical role in prooxidant and anti-oxidant role of p53 [22]. Although their exact roles in salivary gland cancer cells cannot be predicted based on the acquired data. Furthermore, GO enrichment analysis predicted apoptotic proteins might affect IL-33 production. IL-33 was previously suggested as a novel biomarker to distinguish different salivary gland tumours [23]. Additionally, IL-33 expression indicates a favourable prognosis in salivary gland tumours [24].

Finally, IL-33 has been shown to regulate tumour microenvironment by recruiting a series of immune cells that affects progression of various types of cancer [25]. Overall, this project shows a promising approach to transform ACC into an immune “hot” tumour. In future, ANXA3 functionalized nanoparticles should be tested in vivo.

## Methods

### 1. 3D salivary gland cancer cell culturing

2D-salivary gland cancer cell culture was done using UM-HACC-2A cells (adenoid cystic carcinoma) (T8326, abm) cultured in optimized salivary gland medium consisted of PriGroIII (TM003, abm), 10% fetal bovine serum (A56695-01, gibco), 2 mM L-glutamine (G275, abm), 0.4 µg/mL hydrocortisone (H0135-1MG, Sigma Aldrich), 20 ng/mL recombinant human epidermal growth factor (Z100139. abm), 5 µg/mL recombinant human insulin (Z101065, abm), 1% Penicillin/Streptomycin solution (G255, abm); UFH2 (adenoid cystic carcinoma) (kind gift from Prof Maria Zajak Kaye, University of Florida), cultured in Dulbecco’s modified eagle’s medium-high glucose (D5796, Sigma Aldrich), 10% fetal bovine serum (A56695-01, gibco), 1% Penicillin/Streptomycin solution (G255, abm); and A-253 (mucoepidermoid carcinoma) (HTB-41, ATCC) cells cultured in Dulbecco’s modified eagle’s medium-low glucose (D6046, Sigma Aldrich), 10% fetal bovine serum (A56695-01, gibco), 1% Penicillin/Streptomycin solution (G255, abm). The 3D-cancer-immune co-culture models were constructed using a combination of applied extracellular matrix-collagen I (G422, abm) and fibronectin human plasma (F0895-2MG, Sigma Aldrich) as extracellular matrix on ibidi plate with 1.5 polymer coverslip (82426, i-plate 24 well black 14mmibiTreat, ibidi, Germany). NK-92 (natural killer) [CRL-2407, ATCC] cells were used as immune cells. Indirect immune cell coculturing was done initially for cell-based mass spectrometry analysis using Cell culture inserts (Transparent PET membrane) 0.4 µm 20 x 25 mm (NUN140640, Thermo Fisher Scientific), Falcon cell culture insert (Transparent PET membrane) 6 well 3.0 µm (353091, in vitro technologies), and Falcon cell culture insert (Transparent PET membrane) 6 well 0.4 µm (353090, in vitro technologies) were used for co-culturing. For the rest of the entire project, direct immune cell coculturing was done using PBMCs (70025, STEMCELL Technologies). The 3D co-culture model composition and method is disclosed and submitted to IP Australia for provisional patent [Reference].

### 2. FFPE sample processing

Four salivary gland condition FFPE slides were used during the project. (i) Adenoid cystic carcinoma of salivary gland (NBP2-30299, Novus Biologicals), Layout: 1X1, Diameter: 4, Thickness: 5 µm, Age: 51 years, Sex: F, Organ: Parotid gland, Pathology: Adenoid Cystic Carcinoma, Tissue Status: Tumor/Cancer, Species: Human; (ii) normal parotid salivary gland (NBP2-30192, Novus Biologicals), Layout: 1X1, Diameter: 4, Thickness: 5 µm, Age: 37 years, Sex: F, Organ: Parotid gland, Tissue Status: Normal, Species: Human; (iii) mucoepidermoid carcinoma of salivary gland (NBP2-30300, Novus Biologicals), Layout: 1X1, Diameter: 4, Thickness: 5 µm, Age: 31 years, Sex: F, Organ: Parotid gland, Pathology: Mucoepidermoid Carcinoma, Tissue Status: Tumor/Cancer, Species: Human; and (iv) pleomorphic adenoma of salivary gland (NBP2-77923, Novus Biologicals), Layout: 1X1, Diameter: 4, Thickness: 5 µm, Age: 63 years, Sex: M, Organ: Parotid gland, Pathology: pleomorphic adenoma, Tissue Status: Tumor, Species: Human. Deparaffinization process initiated with FFPE samples kept in 60 °C in oven for 1 hour, followed by immersing in xylene two-times for 10 minutes in each round, then 100% ethyl alcohol for 10 minutes, then 95% ethyl alcohol for 10 minutes, then 70% ethyl alcohol for 10 minutes, and finally rehydrated in distill water for 5 minutes.

### 3. H&E staining

Following deparaffinization, dehydration, and rehydration, FFPE samples were immersed in Harris Hematoxylin solution (HHS16-500ML, Sigma Aldrich) for 10 minutes. The slides were washed in running water for 5 minutes. Then, the slides were immersed in Scott’s tap water substitute concentrate (S5134-6X 100ML, Sigma Aldrich) for 60 seconds. The slides were rinsed once in running tap water. Then, the slides were immersed in eosin Y (HT110116-500ML, Sigma Aldrich) for 1 minute. Subsequently, the slides were immersed in 95% ethyl alcohol for 30 seconds, then 100% ethyl alcohol for 2 minutes for 3-times, and then in xylene for 2 minutes for 3-times. Then, Entellan rapid mounting medium (1079610100, Sigma Aldrich) was added onto the samples and coverslip placed. The slides were kept in 4 °C overnight and then transferred to microscopy unit for image acquisition using Olympus_APEXVIEW-APX100 All-in-One Microscope.

### 4. Mass spectrometry sample preparation

Cell and tissue lysates were acquired using sodium deoxycholate triethylammonium bicarbonate (1% in 100mM) (Sigma Aldrich). The lysates were sonicated (short cycle sonication) for 5-10 seconds and then denatured at 95°C for 5 minutes in heating block. The protein amount in each sample (lysates) was determined using Pierce BCA protein assay kit (Thermo Fischer Scientific, Cat 23225). Samples (each 100µg) added to 1% of 1M dithiothreitol (DTT) (Sigma Aldrich, Cat D0632-5G) for 30 minutes at 60°C. Then 4% of 0.5M iodoacetamide (IAA) (Sigma Aldrich, Cat 16125-25GM) was added to the sample and incubated for 30 minutes at room temperature. Then, sequencing grade trypsin (Sigma Aldrich, Cat T6567-20UG) added to the samples and incubated at room temperature overnight. On the subsequent day, the samples were centrifuged at 14000g for 5 minutes. The acquired supernatant was vacuum centrifuged. Finally, 1 µg processed protein sample was added to 0.1% formic acid followed by mass spectrometry using Triple TOF 6600 mass spectrometer (SCIEX). The samples were run following the recommended parameters of Triple TOF 6600-1 hour with 100 variable windows. MS1 accumulation for 0.05 seconds and MS2 accumulation for 0.03 seconds used in the SWATH method ranging from 350-1200 m/z.

### 5. High-pH and reversed phase HPLC

The cancer samples were pooled for the construction of ion library. Two micrograms of each sample were pooled, and fractionated using High pH reverse phase-High performance liquid chromatography. The pooled sample was first vacuum dried then resuspended in mobile phase buffer A (5 mM ammonium hydroxide solution [pH 10.5]). After sample loading and washing with 3% buffer B (5 mM ammonia solution with 90% Acetonitrile [pH 10.5]) for 10 minutes at a flow rate of 300 μL/min, the buffer B concentration was increased from 3% to 30% over 55 minutes, and then to 70% between 55 to 65 minutes and to 90% between 65-70 minutes.

### 6. 2D ion library construction

SWATH_runs (raw files) and 2D information dependent acquisition (IDA) ion library_runs (raw files) were acquired from the Triple TOF 6600 mass spectrometer (SCIEX). ProteinPilot 5.0 (SCIEX) was used for protein identification. Group file(s) were converted to text file(s) using PeakView. All text files (1D-IDAs and 2D-IDAs) were zipped. The seed library (1D-IDA text file) was merged with the 2D-IDA libraries using SwathLibraryMerge on GenePattern.

### 7. SWATH processing

The ion library was imported after quantitation followed by SWATH processing. The multiple comparison of different samples was done using GenePattern server (SwathPairsAndOverall) (Australian Proteome Analysis Facility, Macquarie University).

### 8. GO enrichment analysis

The database for annotation, visualization, and integrated discovery (DAVID) https://david.ncifcrf.gov/ was used for GO analysis. For gene enrichment analysis, all query genes were first converted to ENSEMBL gene ID or STRING database protein IDs. FDR is calculated based on nominal p-value from the hypergeometric test. Fold enrichment was done after dividing the affected protein genes belonging to the pathway, to the corresponding percentage in the background. Only pathways that were in the specified size limits were used for enrichment analysis. After the analysis was done, pathways were first filtered based on the specified FDR cut off. Then the significant pathways were sorted by FDR, fold enrichment, and other metrics. Significant pathways were sorted by average ranks based on FDR (first top pathways were selected by FDR) and fold enrichment. Removal of redundant similar pathways that contained 95% similar genes, more than 50% similar words in their names, and longer pathway names that share similar first 80 characters was done. Assuming protein expressions correspond to respective gene expressions, we additionally compared gene characteristics of proliferation mechanism, immune proteins, and chromosome positioning analysis of the same.

### 9. IHC

Following deparaffinization, dehydration, and rehydration of FFPE samples, citrate buffer (C999-100ML, Sigma Aldrich) 10 mM at pH 6.0 was added and heated at 95 °C – 100 °C for 40 minutes for antigen retrieval. The samples were removed and let cool in the buffer for 30 minutes in room temperature and rinsed with 1X PBS to remove citrate. Then, the FFPE samples were treated twice with 0.5% Triton X-100 in 1X PBS for 30 minutes at room temperature. The samples were blocked with 5% (w/v) bovine serum albumin (A7906-10G, Sigma Aldrich) for 1 hour. Further, the samples were kept at room temperature for drying and liquid repellant slide marker pen (advanced PAP pen, Z377821-1EA, Sigma Aldrich) was used to confine the tissue area on the slides. Subsequently, the FFPE slides were incubated overnight at 4 °C with rabbit anti-cMyb antibody (A304-135A-T, Thermo Fisher Scientific) at a concentration of 1:1000 added to 5% (w/v) bovine serum albumin. Following primary antibody incubation, the slides were washed three times with 1X PBS and then Alexa Fluor 488 Goat anti-rabbit (A32731, Thermo Fisher Scientific) at a concentration of 1:1000 in 5% (w/v) bovine serum albumin added to the FFPE samples for 2 hours at 4 °C and then the samples were fixed and nucleus counter stained with prolong glass with nucblue (P36983, Thermo Fisher Scientific).

### 10. Functionalization of nanoparticles

Gold nanoparticles (40 nm, 20 OD) conjugated to anti-cMyb antibody following manufacturer instruction of gold conjugation kit (Ab154873, abcam). The functionalized gold nanoparticles were kept in 4 °C until further usage. The supplied 5 mg streptavidin coated Poly(lactic-co-glycolic acid) (PLGA) nanoparticles, 120 nm, (CDPLN-10, CD Bioparticles) was reconstituted using 1 mL of DEPC-treated water (BUF kit) (AM9915G, Thermo Fisher Scientific), that made a stock concentration of 5 mg/mL. For antibody-nanoparticle conjugation, 250 µg of streptavidin coated PLGA nanoparticles were added to 5 µg of biotinylated anti-cMyb antibody (biotinylation done following manufacturer instruction of biotin (type B) conjugation mix (ab201796, biotin conjugation kit type b - lightning link, abcam).

### 11. Characterization of functionalized nanoparticles

Optical properties of functionalized gold and PLGA nanoparticles were assessed using UV-visible spectroscopy. Spectra were acquired over a wavelength range of 200-900 nm using JASCO V-760 UV-vis spectrometer (Jasco, Hachioji Japan). The measurements were acquired using 1000 µL of the sample placed in a plastic disposable semi-micro cuvette (Brand Tech, USA). The morphology of functionalized gold and PLGA nanoparticles were assessed after acquiring transmission electron microscopy images (JEOL 1400 TEM, Tokyo, Japan). The surface charge and size (dynamic light scattering) of the nanoparticles were determined using Zetasizer ZS (Malvern Panalytica, Malvern, U.K). Phenom Desktop Scanning Electron Microscope (SEM) with integrated Energy Dispersive X-ray Spectroscopy (EDX) (Thermo Fisher Scientific) was used for elemental analysis of the nanoparticles. Assessment of amide linkage to determine functionalization of nanoparticles were done using iN10 FT-IR microscope (Thermo Fisher Scientific).

Two-dimensional Raman mapping of adenoid cystic carcinoma (ACC) and normal parotid gland FFPE tissue sections treated with anti-cMyb antibody–functionalized gold nanoparticles, together with control samples, was performed using a WITec confocal Raman microscope. A 250 × 250 µm area was scanned using a 633 nm excitation laser with 20 mW power at the sample, 1 s integration time, and a 20× objective. The map consisted of 25 × 25 spectra, corresponding to a step size of 10.4 µm in both x and y directions and an in-plane spatial resolution of approximately 10 µm per pixel.

Raman spectra were pre-processed prior to analysis. Spectra were smoothed using a Savitzky–Golay filter (10-point window) and baseline corrected using the asymmetric least squares (ALS) method. The intensity of the 1650 cm⁻¹ band (amide I region) was used as an indicator of plasmonic nanoparticle presence and local signal enhancement.

For each pixel, the peak intensity at 1650 cm⁻¹ was extracted to generate a two-dimensional intensity map. Image-based intensity quantification and visualization were performed using ImageJ. Raman intensity maps were imported as 8-bit grayscale images, and the full image area was analysed without additional spatial masking. Pixel intensity values were measured using the integrated density/mean grey value metrics. The resulting intensity maps were pseudo-coloured and overlaid onto corresponding bright-field optical images for spatial correlation of nanoparticle localization with tissue morphology.

All samples were processed using identical acquisition parameters and analysis settings to allow direct comparison between ACC, normal tissue, and control conditions.

To determine functionalization of PLGA nanoparticles, ACC and normal parotid gland FFPE samples, and UM-HACC 2A, UFH2, SCC9 cells were incubated overnight at 4 °C with anti-cMyb antibody functionalized PLGA nanoparticles. Following functionalized PLGA nanoparticles incubation, the slides were washed three times with distill water and then Alexa Fluor 488 goat anti-rabbit (2 µg/mL) (A32731, Thermo Fisher Scientific) at a concentration of 1:1000 in distill water added to the FFPE samples and incubated for 2 hours at 4 °C. The samples were fixed and nucleus counter stained with prolong glass with nucblue (P36983, Thermo Fisher Scientific).

Images were acquired using Olympus FV3000 inverted confocal microscope (Olympus, Tokyo, Japan) with 60X silicon oil objective (NA 1.3). Solid status diode laser lines 405nm and 488nm were used for excitation. The reflected signal was directed through a DM 405/488 dichroic mirror and detected within the 430–470 nm range for the DAPI channel or 500–600 nm for the Alexa Fluor 488 channel.

### 12. Overexpression of ANXA3

ANXA3 overexpression was done using Annexin A3 (NM_005139) human recombinant protein (TP301540, OriGene). Streptavidin coated PLGA nanoparticles (250 µg) were added to the mixture of 2.5 µg of biotinylated ANXA3 recombinant protein and 2.5 µg of biotinylated anti-cMyb antibody (biotinylation done following manufacturer instruction of biotin (type B) conjugation mix (ab201796, biotin conjugation kit type b- lightning link, abcam). Non-specific overexpression was done using GAPDH (NM_002046) Human Recombinant Protein (TP302309, OriGene).

### 13. Western blot

UM-HACC 2A, UFH2, UM-HACC-2A_PLGA_ANXA3, UFH2_PLGA_ANXA3, UM-HACC-2A_PLGA_anti-cMyb_ANXA3, and UFH2_PLGA_anti-cMyb_ANXA3, cell lysates were acquired using lysis buffer (R&D Systems, Cat RDSARY009) supplemented with protease inhibitor cocktail (P8340-1ML, Sigma Aldrich) at 1:100 dilution. The lysates were sonicated (short cycle sonication) for 5-10 seconds and then denatured at 95°C for 5 minutes in heating block. The protein amount in each sample (lysates) was determined using Pierce BCA protein assay kit (Thermo Fischer Scientific, Cat 23225). UM-HACC 2A and UFH2 acted as a control, ANXA3 recombinant protein acted as a positive control and PLGA acted as a negative control. Positive control 4 µg, control 10 µg, and the rest of the samples 8 µg was used for western blotting. Pre-stained protein ladder (ab116029, abcam), Rabbit annexin A3 polyclonal antibody (PA5-82483, Thermo Fisher Scientific) at 0.4 µg/mL, Goat anti-Rabbit IgG (H+L) Secondary Antibody, HRP (31460, Thermo Fisher Scientific) at 1:10,000, mouse GAPDH monoclonal antibody ZG003 (398600, Thermo Fisher Scientific) at 2 µg/mL, and Goat anti mouse IgG HRP (31430, Thermo Fisher Scientific) at 1:5000 was used for western blotting. High-resolution images were acquired using Chemidoc MP imaging system and processed using Image Lab version 6.0 (Bio-Rad Laboratories, Hercules, CA, USA).

### 14. 3D spheroid culturing

Spheroid microplate, 96-well polystyrene black with clear round bottom, ultra-low attachment (4520, Corning) with lid was used to create spheroids of adenoid cystic carcinoma cells (10,000 cells per spheroid). ACC spheroids were made using UM-HACC 2A and UFH2 cells at a concentration of 10,000 cells per well of the spheroid microplate was incubated at humidified 37 °C and 5% CO_2_ for 24 hours.

### 15. Activation of CD4+/CD8+ lymphocytes

Human PBMC 1.5 x 10^7^ cells/vial (70025, STEMCELL Technologies) was used for direct spheroid immune coculturing assays. For activating 7.5 x 10^6^ PBMCs, 150 µL of CD3/CD28 magnetic beads stock solution with a concentration 5 x 10^4^ beads/µL was used in RPMI 1640 medium supplemented with 10% FBS, 1% Penicillin/Streptomycin. Control experiment was done using non-activated 7.5 x 10^6^ PBMCs in RPMI 1640 medium supplemented with 10% FBS, 1% Penicillin/Streptomycin. Both activated and non-activated PBMCs were incubated at humidified 37 °C and 5%CO_2_ for 24 hours. After incubation, the magnetic beads were extirpated using magnetic stand (MB-02, Acro Biosystems).

### 16. Determination of immune infiltration

Both activated and non-activated PBMCs were added equally (approximately 3 x 10^5^ cells) to UM-HACC 2A spheroids on ECM, UFH2 spheroids on ECM, anti-cMyb antibody functionalized PLGA nanoparticles conjugated to ANXA3 recombinant protein treated UM-HACC-2A spheroids on ECM, and anti-cMyb antibody functionalized PLGA nanoparticles conjugated to ANXA3 recombinant protein treated UFH2 spheroids on ECM, on each well of ibidi plate with 1.5 polymer coverslip (82426, i-plate 24 well black 14mmibiTreat, ibidi, Germany).

After incubation, each ACC spheroid immune coculture on ECM (ANXA3 overexpressed or untreated) were fixed using 4% paraformaldehyde and further incubated at 4 °C overnight. Subsequently, ECM was permeabilized using 0.5% Triton X-100 in 1X PBS added to each well and incubated at room temperature for 1 hour. Anti-CD8 alpha antibody [CAL66] (AB237709-1001, abcam) (1:100), Anti-CD4 antibody [EPR6855] – Mouse IgG1 (Chimeric) (AB317787-1007, abcam) (1:100), Alexa Fluor 568 goat anti-mouse IgG (H+L) (A11004, Thermo Fisher Scientific) (2 µg/mL), Alexa Fluor 488 Goat anti-rabbit (A32731, Thermo Fisher Scientific) (2 µg/mL), and prolong glass with nucblue (P36983, Thermo Fisher Scientific) was used for staining CD4+ and CD8+ lymphocytes. Quantification of the immune infiltration was based on measuring fluorescence intensity using FV31S-SW software (Version: 2.6.1.243, OLYMPUS CORPORATION, Japan).

### 17. Size of spheroids

Spheroid immune cell cocultures established on the extracellular matrix (ECM) model were imaged using an APX100 benchtop fluorescence microscope (Olympus, Tokyo, Japan). Spheroid size was quantified using ImageJ software (National Institutes of Health, Bethesda, MD, USA).

### 18. Apoptotic profiler

Apoptotic profiler samples were prepared from the acquired different treatment conditions from the coculture models. The adenoid cystic carcinoma cells were added to 100 μL of lysis solution, supplied in Proteome Profiler Human Apoptosis Array Kit (R&D Systems, Cat RDSARY009) and 10 μL of protease inhibitor (Protease Inhibitor Cocktail I, Tocris, Cat 5500). The lysates were stored in -80°C freezer for further assays. Upon following manufacturer instruction of Proteome Profiler Human Apoptosis Array Kit (R&D Systems, Cat RDSARY009), high-resolution images were acquired using Chemidoc MP imaging system and processed using Image Lab version 6.0 (Bio-Rad Laboratories, Hercules, CA, USA). Quantification of relative protein expressions were done using microarray profile plugin in ImageJ (ImageJ 1.52a, National Institutes of Health, USA).

### 19. Statistical Analysis

Multiple comparisons of the generated SWATH output were performed using GenePattern server. ANOVA p-values were adjusted for multiple testing by Benjamini–Hochberg false discovery rate adjustment. Unpaired t-test was done for each pairwise comparison to determine significant upregulated and downregulated proteins. Fold change cut-off 1.5 and p-value cut-off 0.05 was used during the entire analysis.

## Author contributions

Conceptualization: RC; Project and experimental design: RC, AC, MS, AT, KS; Funding Acquisition: RC, KS; Methods: RC, RS, AC, AA, TW, MS, AT, KS; Data acquisition: RC, RS, AC, MA, CS, AA, TW, AT; Statistical Analysis: AC, MA, CS, AA, TW, AT; Manuscript write up: RC; Manuscript corrections: RS, AC, CS, AA, MS, AT, KS; Supervision:. KS

## Ethics

The project was done after Macquarie University Medical Sciences Human Research Ethics Committee approval to use FFPE samples “19710 - Chakraborty-A Prelude towards transforming adenoid cystic carcinoma into an immune hot tumour”. Cell and spheroid immune coculture work was done upon acquiring biosafety approval from Institutional Biosafety Committee, Macquarie University, “A tumour-cell based initiation of immune evasion – 16835”.

## Acknowledgement

The project is funded by the Adenoid Cystic Carcinoma Research Foundation (ACCRF, Massachusetts, USA) and Australian and New Zealand Head and Neck Cancer Society (ANZHNCS, Australia). We are grateful to Jeffrey Kauffman (Co-founder of ACCRF) and Dr Nicole Spardy Burr (Executive Director, ACCRF) for their immense help towards the project and providing me useful information and suggestions regarding experimental design of the project. We are grateful to the scientific committee of ANZHNCS for their continuous trust and support towards adenoid cystic carcinoma research. We would like to acknowledge the support from Prof Yuling Wang group at the School of Natural Sciences, Macquarie University, and the staff of microscopy unit and Australian Proteome Analysis Facility, Faculty of Science and Engineering, Macquarie Analytical and Fabrication Facility (MAFF). Macquarie University. We would like to acknowledge the support of Prof Jacques Nor (University of Michigan), Prof Fredrick Kaye (University of Florida), Prof Fiona Simpson (University of Queensland), and Dr Charbel Darido (University of Melbourne) for their initial technical support during the culturing of adenoid cystic carcinoma cells.

## Funding information

The project is funded by Adenoid Cystic Carcinoma Research Foundation (Massachusetts, USA) and Australian and New Zealand Head and Neck Cancer Society Trudi Shine Adenoid Cystic Carcinoma Grant Award 2025

## Data Availability

The datasets generated and analysed during the current study are available in the figshare repository, https://doi.org/10.6084/m9.figshare.25990132

## Conflict of Interest

The authors declare no competing interests

## Notes

### Competing Interest Statement

The authors have declared no competing interest.

