## Supplementary for "Functionalized nanoparticle transforms “cold” to “hot” adenoid cystic carcinoma of salivary gland tumour microenvironment in vitro"

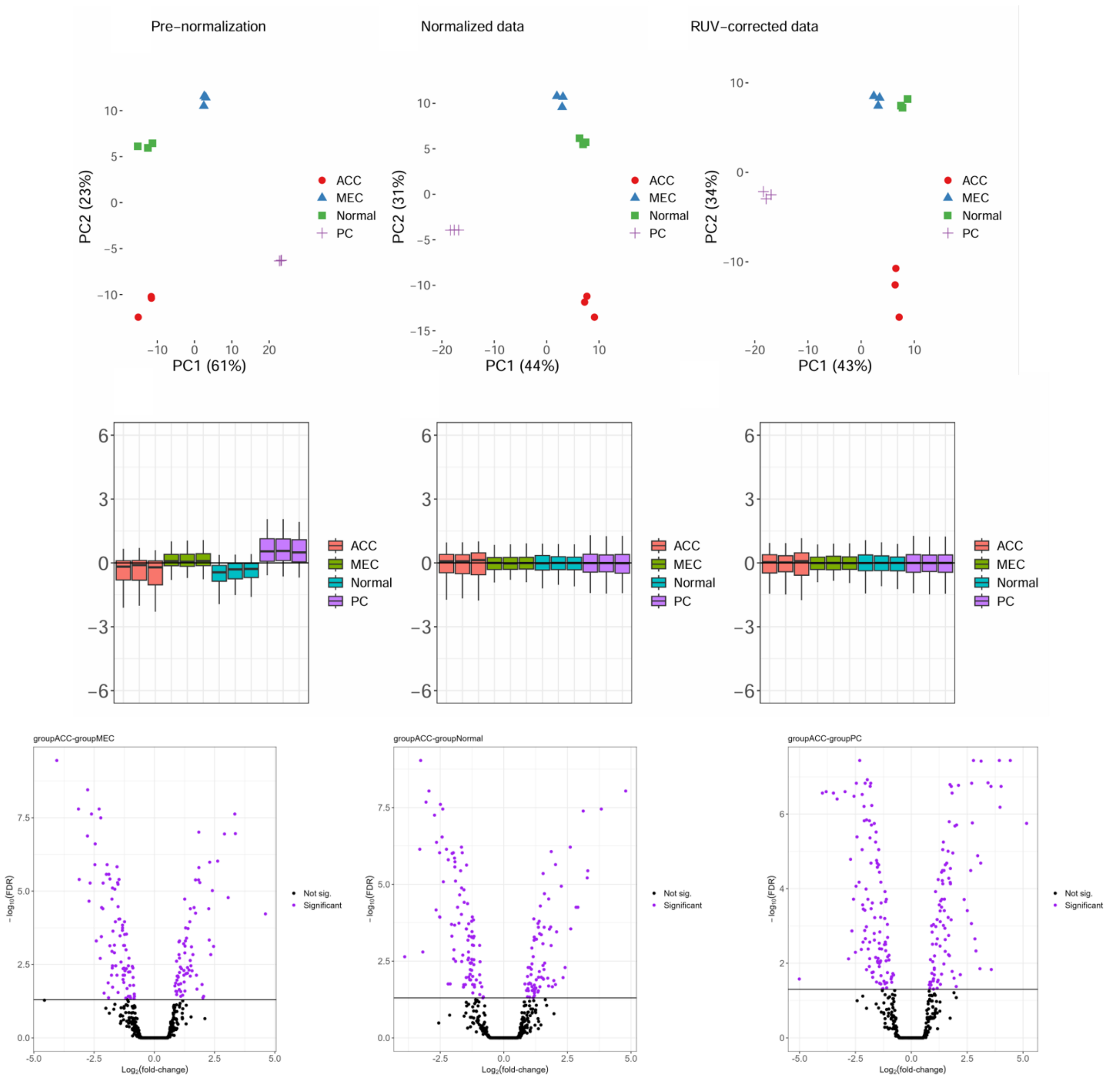


**Supplementary Figure 1. Mass Spectrometry Analysis of salivary gland tissues.** Principal component analysis and volcano plots of the mass spectrometry samples of pleomorphic adenoma of salivary gland, mucoepidermoid carcinoma of salivary gland, adenoid cystic carcinoma of salivary gland, and normal parotid salivary gland.

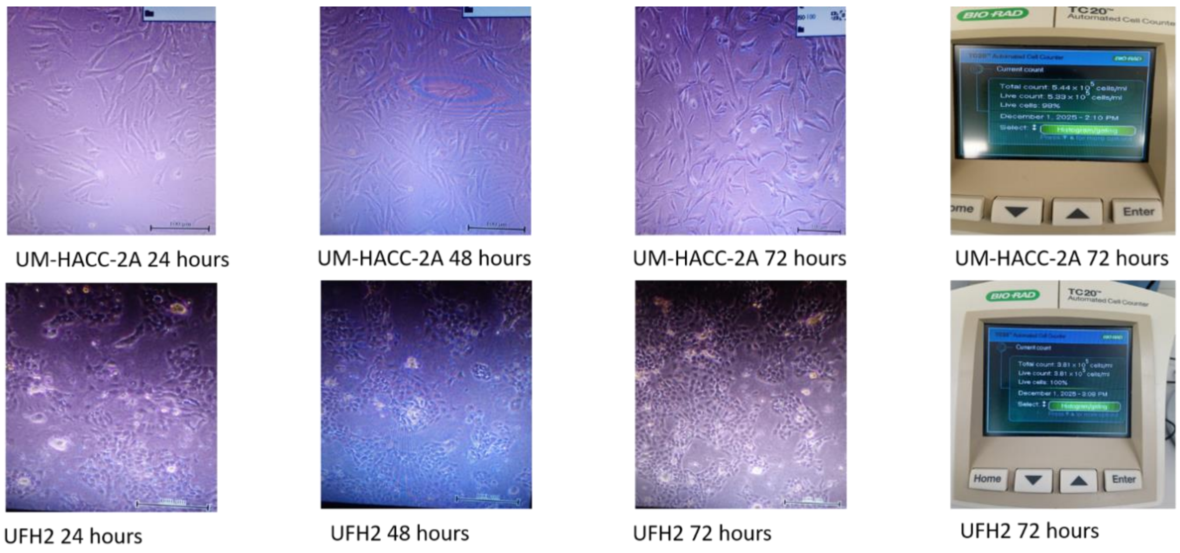


**Supplementary Figure 2. Morphology of cells on ECM.** UM-HACC-2A and UFH2 cells on ECM for 24 hours, 48 hours, and 72 hours. Trypan blue assay showing high percentage of live cells. Images acquired with 4X objective. Scale 200 µm.

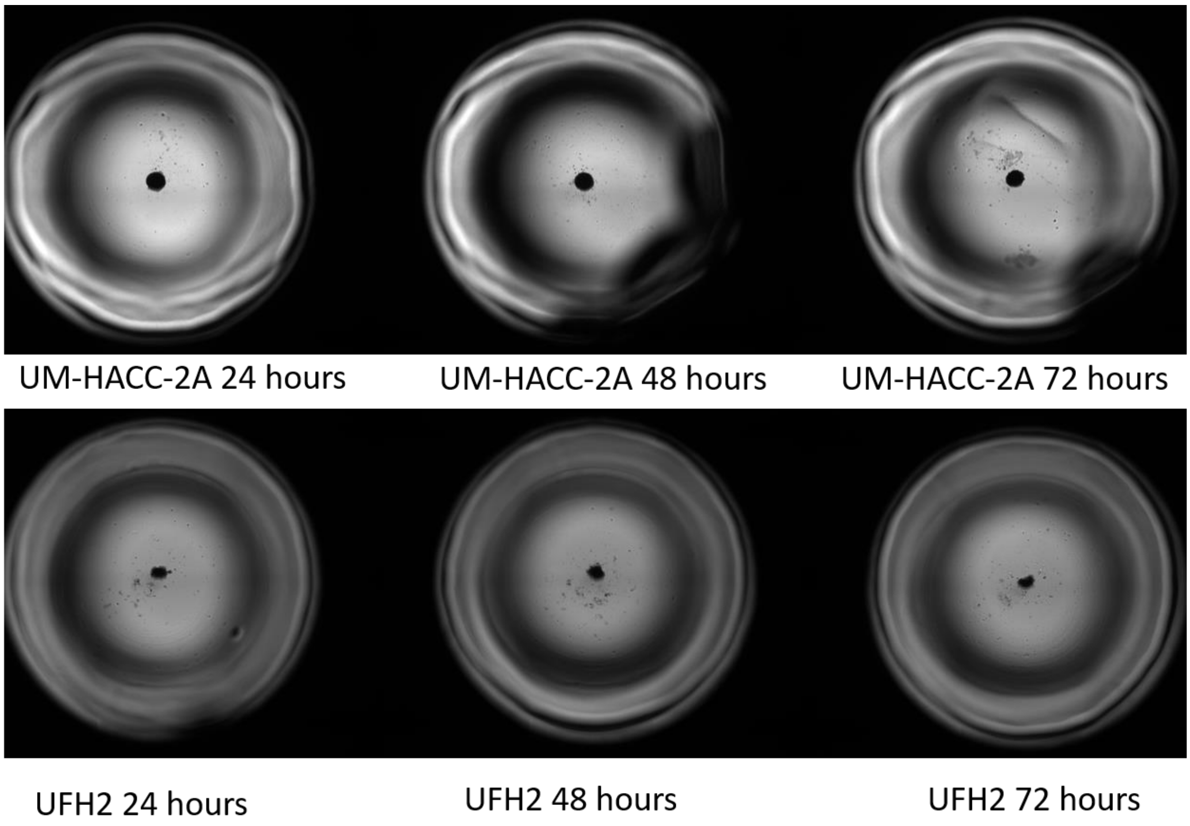


**Supplementary Figure 3. Morphology of spheroids on ECM.** UM-HACC-2A and UFH2 spheroids on ECM for 24 hours, 48 hours, and 72 hours. Wide field images acquired with 4X objective. Scale 200 µm.

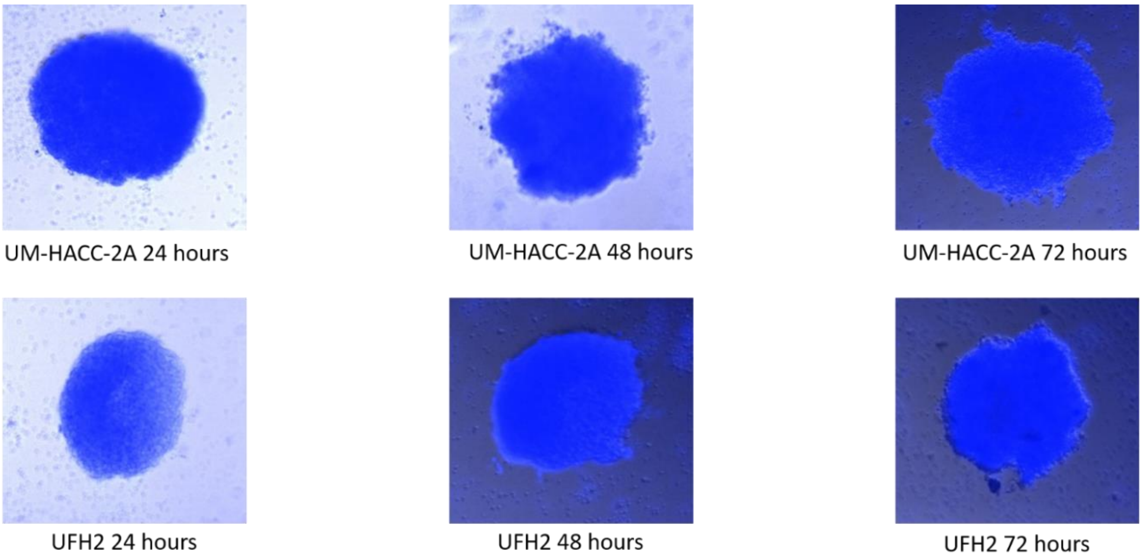


**Supplementary Figure 4. Morphology of spheroids on ECM in PBMC coculture.** UM-HACC-2A and UFH2 spheroids on ECM in PBMC coculture for 24 hours, 48 hours, and 72 hours. Wide field images acquired with 4X objective. Scale 200 µm.

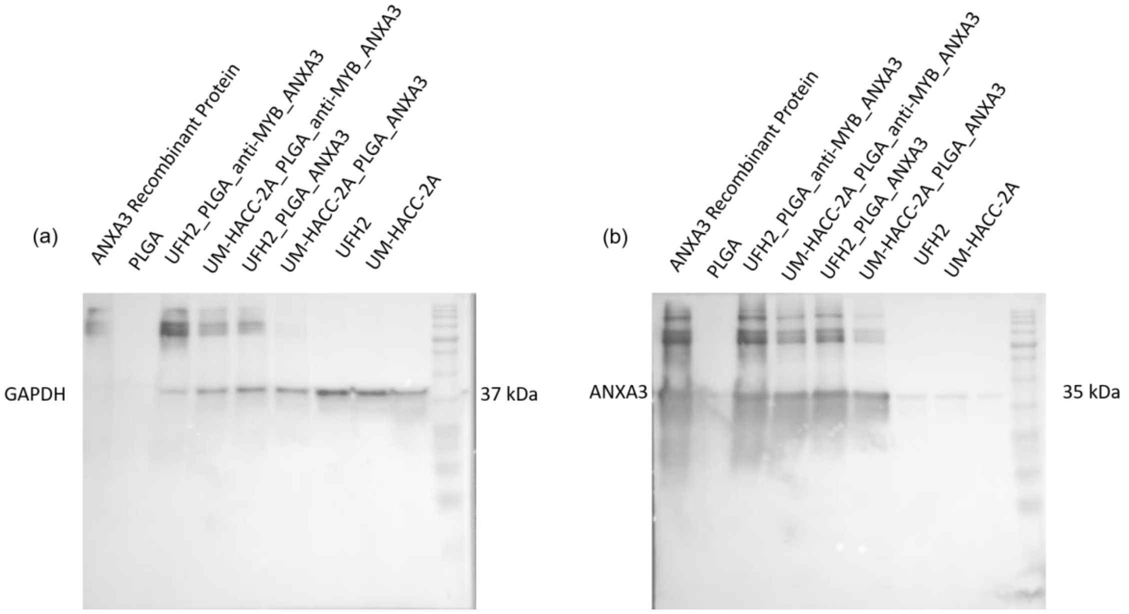


**Supplementary Figure 5. Western blot (full blot) showing overexpression of ANXA3 using anti-cMyb antibody-ANXA3-PLGA nanoparticles.** UM-HACC 2A and UFH2 acted as a control, ANXA3 recombinant protein acted as a positive control and PLGA acted as a negative control. Positive control 4 µg, control 10 µg, and the rest of the samples 8 µg was loaded. Pre-stained protein ladder (ab116029, abcam), Rabbit annexin A3 polyclonal antibody (PA5-82483, Thermo Fisher Scientific) at 0.4 µg/mL, Goat anti-Rabbit IgG (H+L) Secondary Antibody, HRP (31460, Thermo Fisher Scientific) at 1:10,000, mouse GAPDH monoclonal antibody ZG003 (398600, Thermo Fisher Scientific) at 2 µg/mL, and Goat anti mouse IgG HRP (31430, Thermo Fisher Scientific) at 1:5000.

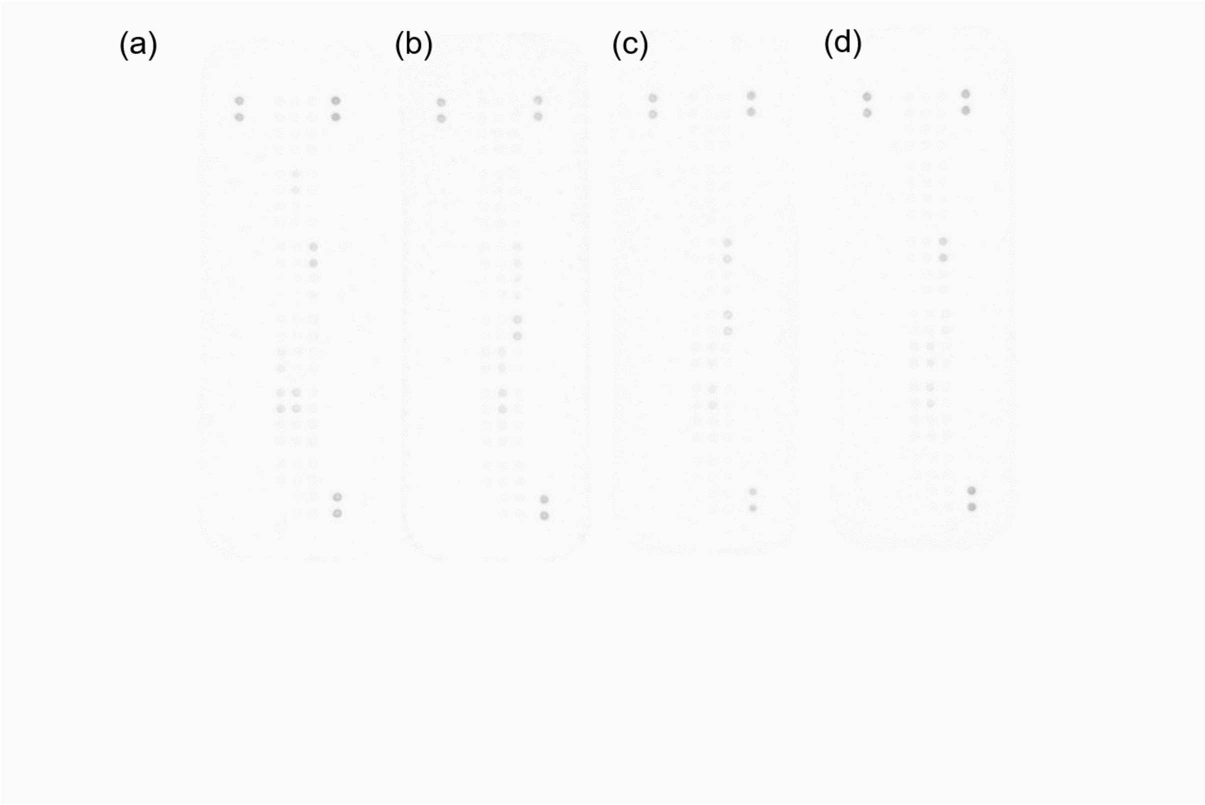


**Supplementary Figure 6. Proteome Profiler Human Apoptosis Array (full blot).** (a) anti-cMyb antibody-ANXA3-PLGA nanoparticles on UM-HACC 2A spheroids in PBMC coculture; (b) UM-HACC 2A spheroids in PBMC coculture; (c) UFH2 spheroids in PBMC coculture; anti-cMyb antibody-ANXA3-PLGA nanoparticles on UM-HACC 2A spheroids in PBMC coculture. Images were acquired using Chemidoc MP imaging system and processed using Image Lab version 6.0 (Bio-Rad Laboratories, Hercules, CA, USA).
